# Discovery of dehydroamino acids and their crosslinks in Tau and other aggregating proteins of Alzheimer’s disease

**DOI:** 10.1101/2024.09.23.614558

**Authors:** Samuel W. Markovich, Brian L. Frey, Mark Scalf, Michael R. Shortreed, Lloyd M. Smith

## Abstract

**Background:** Alzheimer’s disease (AD) is characterized by accumulation of two types of protein aggregates, senile plaques and neurofibrillary tangles, which strongly contribute to the pathogenesis of the disease. Senile plaques consist primarily of aggregated amyloid-ß, while neurofibrillary tangles form via aggregation of the protein Tau, as well as other microtubule-associated proteins such as CRMP2. Posttranslational modifications have been hypothesized to contribute to the initial aggregation events that lead to SPs and NFTs. Dehydroamino acids (DHAAs) are posttranslational modifications rarely observed in humans and have not previously been reported in AD. DHAAs arise from the eliminylation of serine, threonine, or cysteine, yielding a double bond with distinct molecular geometry and reactivity. Their geometry can produce secondary structure rearrangements, such as those seen in senile plaque and neurofibrillary tangle formation, while their reactivity can cause intramolecular or intermolecular (protein-protein) crosslinking. We hypothesized that this modification might be present in protein aggregation-associated neurodegenerative disorders like AD.

**Methods:** We performed mass spectrometry-based bottom-up proteomics on the sarkosyl-insoluble (protein aggregate-enriched) material from ten AD brains and three age-matched controls. Identifications of DHAA-mediated crosslinked peptides were validated using both an isotopic labeling strategy and spike-in experiments employing synthetic crosslinked peptide standards. Similar findings were obtained in searches of publicly available proteomic datasets from AD and control brains.

**Results:** We identified 412 sites of DHAA modification in 184 proteins, with the highest prevalence in the neurofibrillary tangle-forming proteins Tau and CRMP2. Comparison with results of previous protein aggregate interactomics studies show that proteins containing the DHAA modification are more highly associated with protein aggregates than are proteins containing any other individual posttranslational modification. We further observed 11 protein crosslinks arising from DHAAs, including three from the Tau protein. Label free quantification showed that Tau crosslinks are an order of magnitude more prevalent in AD samples than in age-matched controls.

**Conclusions:** Dehydroamino acids and their derivatives are prevalent modifications in the Alzheimer’s disease brain proteome. These modifications give rise to protein crosslinks which may contribute to protein aggregation processes.

## Background

Alzheimer’s disease (AD) is a widespread neurodegenerative disorder that accounts for ∼70% of all cases of dementia [1]. A hallmark of AD is the formation of two distinct protein aggregates: senile plaques (SPs) and neurofibrillary tangles (NFTs), which strongly contribute to the pathogenesis of the disease. SPs consist primarily of aggregated amyloid-β, while NFTs commonly form via aggregation of the protein Tau, as well as other microtubule-associated proteins such as CRMP2 [2,3]. The spread of SPs and NFTs has been thoroughly investigated, yet it remains unclear how the constituent proteins begin aggregating [4]. While posttranslational modifications (PTMs) such as phosphorylation have been strongly implicated [2,5], more comprehensive knowledge of the nature, prevalence, and possible roles of the PTMs present in aggregating proteins could greatly benefit the understanding of AD and other protein aggregation disorders.

Dehydroamino acids (DHAAs) are noncanonical amino acids that form posttranslationally in proteins through eliminylation [6]. Eliminylation is a particular form of β-elimination that occurs via (a) the dehydration of serine or threonine, or (b) the loss of hydrogen sulfide from cysteine, or (c) the elimination of phosphate from serine and threonine, or thiophosphate from cysteine. Eliminylation is known to occur more readily from phosphorylated precursors, and we thus hypothesized that hyperphosphorylated proteins such as Tau may contain this modification [7]. DHAAs contain electrophilic alkenes that can spontaneously form crosslinks through Michael addition with nucleophilic amino acid residues, such as cysteine, lysine, or histidine [8]. The formation and subsequent chemical reactions of DHAAs are summarized in Fig. 1. In addition to their ability to form crosslinks, they can react with cellular nucleophiles such as glutathione or homocysteine [8]. Interestingly, DHAAs have also been shown to have the capacity to alter the secondary structure of model peptides without crosslinking [9].

**Fig. 1:**
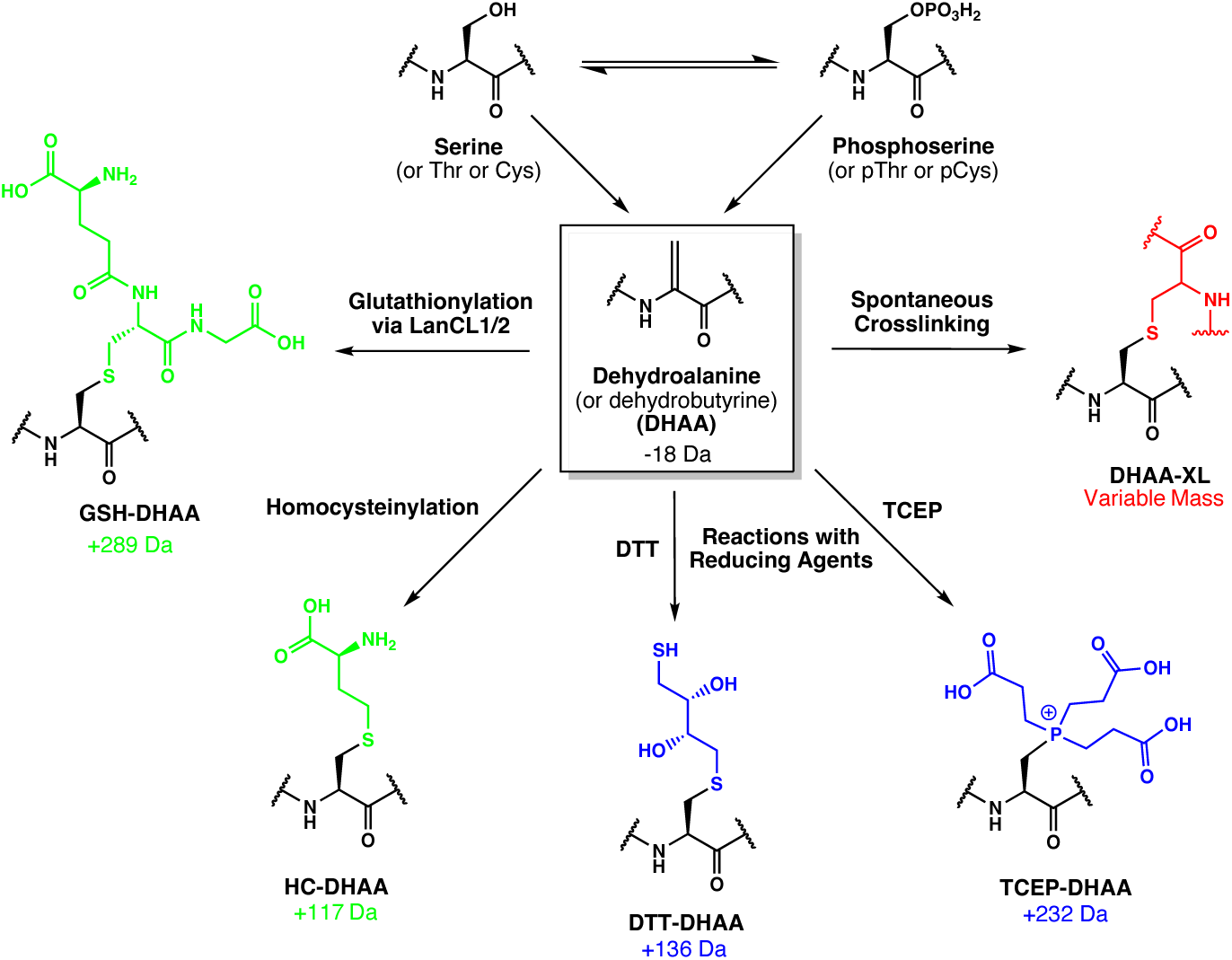
Dehydroamino acids and their conjugates. Serine or cysteine residues within proteins may be eliminylated to form dehydroalanine, while threonine eliminylation yields dehydrobutyrine; these modified residues are dehydroamino acids (DHAAs) that contain an electrophilic alkene. DHAAs can react with common biological nucleophiles such as glutathione (GSH) and homocysteine (HC) to form the corresponding conjugates (green), or with nucleophilic side chains (e.g. cysteine, histidine, lysine) in protein molecules to form intramolecular or intermolecular crosslinks (DHAA-XL) (red). They may also react *in vitro* with nucleophilic reducing agents such as dithiothreitol (DTT) or tris(2-carboxyethyl)phosphine (TCEP) that are often added to samples in proteomic workflows (blue). The approximate mass shifts corresponding to each of these modifications are shown.

DHAAs have been most commonly observed in microorganisms, where they are formed enzymatically in small molecules or within proteins [7]; in contrast, they have been observed only rarely in human proteins. The evidence for DHAAs is weak in some of the human examples [10–12], as the loss of H_2_O/H_2_S (−18/-34 Da) may be generated artifactually during mass spectrometry (MS) [13]. Stronger evidence is obtained from the reaction of DHAAs *in vivo* or *in vitro* with a nucleophile, yielding what we refer to here as DHAA conjugates. These conjugates allow us to infer the presence of an eliminylation site from MS observation of their unique mass shifts. DHAA conjugates were identified in the capsid and matrix proteins of HIV virions grown in a human cell line [14], human kinases [15], and αB-crystallin from human lens tissue [8,16,17]. Although the origin of these modifications in humans is unknown, authors commonly suggest that DHAAs are the result of oxidative damage, as there is no known enzyme in the human proteome capable of catalyzing eliminylation (though some homologues to known DHAA-generating enzymes have been suggested) [17–19].

The most thorough characterization of DHAA-related species in human samples to date occurred in the study of αB-crystallin, isolated from human lens tissue [8]. The authors discovered two species that were formed through DHAA intermediates: dehydroamino acid-mediated crosslinks (DHAA-XLs), and glutathionylated dehydroamino acids (GSH-DHAAs). The crosslinks were formed by reaction of the DHAAs with nucleophilic cysteine or lysine residues also present within the αB-crystallin; and the glutathione modifications formed by reaction of the DHAAs with glutathione. Notably, αB-crystallin is known to aggregate in the formation of cataracts. The authors hypothesized that DHAA-XLs contributed to the αB-crystallin aggregation, and that glutathionylation of DHAAs plays a protective role by blocking the formation of DHAA-XLs, thereby preventing DHAA-mediated protein aggregation.

The authors of the αB-crystallin study suggested that glutathionylation of DHAAs occurs spontaneously by reaction with free, intracellular glutathione; however, more recent work indicates a possible enzymatic mechanism [15]. Lai et al. recently showed that a family of proteins expressed in humans known as the lanthionine synthase C-like proteins (LanCLs) catalyze the addition of glutathione to DHAAs, providing an enzymatic route to formation of GSH-DHAAs. Interestingly, LanCL1 has been shown to be vital for neuronal development and has been implicated in mitigating oxidative stress in the brains of mouse models [20]. It has also been shown that LanCL1 overexpression promotes neuron survival in ALS mouse models [21], and it is naturally upregulated in the brains of mice treated with MPTP, a parkinsonian neurotoxin [22]. These indications of an important role for LanCL1 in murine brain model systems suggest that DHAAs are present in mammalian brain samples and that glutathionylation of DHAAs has a neuroprotective role.

Herein, we report the first observations of DHAAs, DHAA conjugates, and DHAA-mediated crosslinks in the protein aggregate-enriched fraction of brain specimens from Alzheimer’s disease and control patients. DHAAs, GSH-DHAAs, homocysteinylated DHAAs (HC-DHAAs), and DHAA-XLs were observed in Tau, as well as in other proteins implicated in AD. We first observed these DHAA-species in publicly available AD proteomic datasets [23], and then reproduced the findings in samples prepared in our laboratory. Their identities were experimentally confirmed using an isotopic labeling strategy that distinguishes crosslinked peptides from peptide monomers [24]. Tau crosslinks were further validated using synthetic standards spiked into brain samples. Label-free quantification indicated that many DHAA species, particularly those found in Tau, are more abundant in AD samples than age-matched controls. These observations, considered together with the literature reports indicating important roles for glutathionylation by LanCL enzymes in neurodegenerative disorders, suggest that DHAA-mediated crosslinking may play an important role in AD pathogenicity.

## Methods

### Research participants and ethics approvals

Angular gyrus brain tissue samples (Brodmann area 39) from ten Alzheimer’s disease and three age-matched control patients were provided by the Wisconsin Brain Donor Program (WBDP) and the Wisconsin Alzheimer’s Disease Research Center (WADRC) (Table S1). Recruitment and informed consent for brain donation is administered under the WBDP and WADRC protocols approved and monitored by the University of Wisconsin-Madison Institutional Review Board in compliance with the Declaration of Helsinki. As part of the consent process, participants or their legal representatives provide written informed consent for brain donation and use of the collected brain tissue for sharing and investigation of brain health and neurodegenerative disease-related research. Requests for tissue samples are reviewed for appropriateness of scope and feasibility by the WBDP resource committee prior to approval and tissue sharing (https://www.adrc.wisc.edu/apply-resources). Since this study used only decedent tissue samples and the scope was consistent with WBDP protocols, this study was determined to meet the standards of minimal risk and was exempted from local IRB oversight after their review.

### Preparation of sarkosyl-insoluble protein from brain tissue samples

Tissue from each sample was cut into approximately 250 mg pieces and frozen in liquid nitrogen for 5 min. Homogenization by pulverization was performed using a Cellcrusher stainless-steel cup and piston unit that had been kept in liquid nitrogen 15 min prior to use. Homogenized tissue was transferred to a 2 mL Eppendorf tube and washed once with phosphate buffered saline. The tissue was spun down at 125x*g*, 4°C for 10 min. Tissue was resuspended in prechilled 1 mL homogenization buffer (25mM Tris-HCl, pH 7.5, 150 mM NaCl, 10 mM EDTA, 10 mM EGTA, 1.2x Thermo Scientific™ Halt™ Protease and Phosphatase Inhibitor Cocktail (Halt PPI)) and kept on ice. Dounce homogenization was used as needed for incompletely homogenized tissue. To each tube, 80 μL prechilled 5 M NaCl and 120 μL prechilled 10% sarkosyl detergent were added for a final concentration of 500 mM NaCl and 1% sarkosyl detergent. Samples were incubated at 4°C for one hour. Tissue was lysed by sonication on ice using a Model 505 Sonic Dismembrator (Fisherbrand) for 3s on, 3s off for 5 cycles. Brain lysis material was clarified by centrifugation at 11,000x*g* for 30 min at 4°C. The supernatant was transferred to a new tube. Each sample was ultracentrifuged at 100,000x*g* for 2 hours at 4°C (Optima MAX-TL). Sarkosyl-soluble sample (supernatant) was transferred to a new tube and brought up to 2% SDS. The sarkosyl-insoluble sample (pellet) was washed with 100μL 18.2MΩ water and redissolved in 200 µL 4% SDS, 50mM Tris-HCl pH 7.5. Protein concentration was determined using Pierce^TM^ 660nm Protein Assay with the ionic detergent compatibility reagent (Thermo Fisher Scientific). Sarkosyl-soluble aliquots of 100 μg and sarkosyl-insoluble aliquots of 40 μg were stored at - 80°C.

### Preparation of bottom-up proteomics samples

Samples were reduced with 5mM Bond-Breaker^TM^ TCEP (Thermo Fisher Scientific) and alkylated with 30mM iodoacetamide (Sigma-Aldrich). Samples were prepared for mass spectrometry using the SP3 solid phase purification method as described [25]. Briefly, Sera-Mag™ carboxylate-modified magnetic beads (500 μg, Cytiva) were added to each 100 μL sample, followed by an equal volume of molecular biology grade absolute ethanol (Thermo Fisher Scientific). Samples were mixed for 5 min at room temperature, followed by brief benchtop centrifugation. Samples were incubated on a magnetic stand for 2 min. The supernatant was removed. Beads were washed by resuspension in 500 μL 80% ethanol, placed on a magnetic stand for 2 min, and the supernatant was removed. This wash was repeated three times for a total of four washes. Beads were resuspended in 100 μL 50 mM Tris-HCl pH 7.5 with 2 μg of trypsin (Promega) (1:20 ratio) and nutated at 47°C for 2 hours. Samples were centrifuged at 20,000x*g* for 1 min at room temperature and carefully transferred to a magnetic stand to not disturb the pelleted beads. The supernatant, containing tryptic peptides, was transferred to a new tube. Samples were immediately desalted using a C18 solid-phase extraction pipette tip (Pierce™ C18 Pipette Tips, 100μL, Thermo Fisher Scientific) according to manufacturer’s protocols. Desalted samples were dried using a SpeedVac Concentrator and reconstituted in 95:5 H_2_O:ACN, 0.1% formic acid for mass spectrometric analysis.

### Preparation of H_2_^18^O-labeled bottom-up proteomics samples

Back et al. described an isotopic tagging strategy to distinguish crosslinked peptides from non-crosslinked peptides based upon the presence of two terminal carboxylates in the crosslinked product, in contrast to the single terminal carboxylate present in a standard linear peptide [24]. Tryptic digestion of a protein mixture in H_2_^18^O leaves ^18^O atoms for both C-terminal carboxylate oxygens. The 2 Da mass shift for each oxygen thus gives rise to an overall 4 Da mass shift for a linear peptide and an overall 8 Da mass shift for a crosslinked pair of peptides. Samples were reduced, alkylated, and prepared for tryptic digestion using the SP3 processing steps described above. Beads were resuspended in 100 μL 50mM Tris-HCl pH 7.5 in H_2_^18^O (95% isotopic purity, Cambridge Isotope Laboratories) containing 2 μg trypsin and nutated at 47°C for 2 hours. Samples were then processed as described above to prepare the tryptic peptides for mass spectrometric analysis.

### High pH-offline fractionation

Desalted peptides were fractionated using high performance liquid chromatography (HPLC) (nanoAcquity, Waters). HPLC separation employed a 365 μm diameter fused silica capillary micro-column packed with 25 cm of 3 μm-diameter, 300 Angstrom-pore-size, C18 beads (Phenomenex, Jupiter). Peptides were loaded on-column at a flowrate of 2 µL/min for 15 min and then eluted over 40 min at a flowrate of 2 µL/min with a gradient of 0% to 70% acetonitrile in 20mM ammonium formate, pH 10.0. Table S2 lists collection times for 16 eluate samples that were pooled into Fractions 1-8. Fractions were dried using a SpeedVac Concentrator and reconstituted in 95:5 H_2_O:ACN, 0.1% formic acid or mass spectrometric analysis.

### Synthesis of crosslinked peptide standards

We synthesized crosslinked peptides as synthetic standards for two DHAA-XLs using the reactions shown in Schemes S1 and S2 (Supplementary Material 1). Barium hydroxide may be used to convert phosphoserine to dehydroalanine (DHA) within peptides [26]. Once a DHA has been generated, it is available to react with free thiols, like those present in cysteine side chains, to form the resulting thioether.[8,26] We installed the DHA modification at the site of the phosphoserine eliminylation in one peptide and reacted that DHA-modified product with the cysteine-containing partner peptide. Custom peptides IG[pS]TENLK, SPVV[pS]GDTSPR, and CGSKDNIK were purchased from Thermo Scientific Custom Peptide Synthesis Service, where [pS] represents phosphoserine. Peptides were dissolved to a final concentration of 2 mg/mL in HPLC-grade water (Fisherbrand) and stored in aliquots at −20°C. 100 μL of 2 mg/mL IG[pS]TENLK was mixed 1:1 with 250 mM Ba(OH)_2_ (Sigma-Aldrich) in a total volume of 200 μL and incubated at 37°C for 90 min. Barium cations were precipitated by addition of 35 μL 1M Na_3_PO_4_ buffer, pH 7.0 (Teknova) with vigorous mixing. Mixtures were clarified by centrifugation at 20,000x*g* for 1 min at room temperature. The supernatant, which contained the eliminylated peptide (IG[DHA]TENLK, where DHA represents dehydroalanine) in approximately 50 mM Na_3_PO_4_, was transferred to a new tube, and the pH was adjusted with 12% H_3_(PO_4_) to pH 10, aliquoted into 25 μg fractions, and dried using a SpeedVac Concentrator. Aliquots were stored at −20°C.

To generate the crosslinked species, 25 μg of IG[DHA]TENLK, was mixed with 50 μg of CGSKDNIK in 100 μL 250mM Na_3_PO_4_, pH 10.0. Samples were incubated at 70°C for 24 hours, yielding the IG<u>[DHA]</u>TENLK-<u>C</u>GSKDNIK crosslinked species, where the underlined residues are those involved in the crosslink. Samples were immediately desalted using a C18 solid-phase extraction pipette tip (Pierce™ C18 Pipette Tips, 100μL, Thermo Fisher Scientific) according to manufacturer’s protocols. Desalted samples were dried using a SpeedVac Concentrator and reconstituted in 95:5 H_2_O:ACN 0.1% formic acid for mass spectrometric analysis. The same protocol was used to generate the SPVV<u>[DHA]</u>GDTSPR-<u>C</u>GSKDNIK crosslinked species.

### High performance liquid chromatography-tandem mass spectrometry (HPLC-MS/MS) analysis

Fractions were analyzed by HPLC-MS/MS using a system consisting of an HPLC (nanoAcquity, Waters) connected to an electrospray ionization (ESI) Orbitrap mass spectrometer (QE-HF, ThermoFisher Scientific). HPLC separation employed a 365 μm diameter fused silica capillary micro-column packed with 20 cm of 1.7μm-diameter, 130 Angstrom-pore-size, C18 beads (Waters BEH), with an emitter tip pulled to approximately 1 μm using a laser puller (Sutter instruments). Peptides were loaded on-column at a flowrate of 400 nL/min for 30 min and then eluted over 120 min at a flowrate of 300 nL/min with a gradient of 5% to 35% acetonitrile, in 0.1% formic acid. Full-mass profile scans were performed in the FT orbitrap between 375-1500 m/z at a resolution of 120,000, followed by MS/MS HCD scans of the ten highest intensity parent ions at 30% relative collision energy and 15,000 resolution, with a mass range starting at 100 m/z. Dynamic exclusion was enabled with a repeat count of one over a duration of 30 seconds. Spike-in experiments utilized an inclusion list for the two species of interest, the IG<u>[DHA]</u>TENLK-<u>C</u>GSKDNIK crosslink (m/z: 474.47400, z=4, 45-75 min) and SPVV<u>[DHA]</u>GDTSPR-<u>C</u>GSKDNIK (m/z:487.49546, z=4, 45-75 min).

### Data analysis

The data analysis process used for the confident identification of DHAA modified residues, as well as their conjugates and crosslinks, from a large quantity of deep proteomic data obtained from a total of 105 brain specimens, is shown in Fig. 2.

**Fig. 2:**
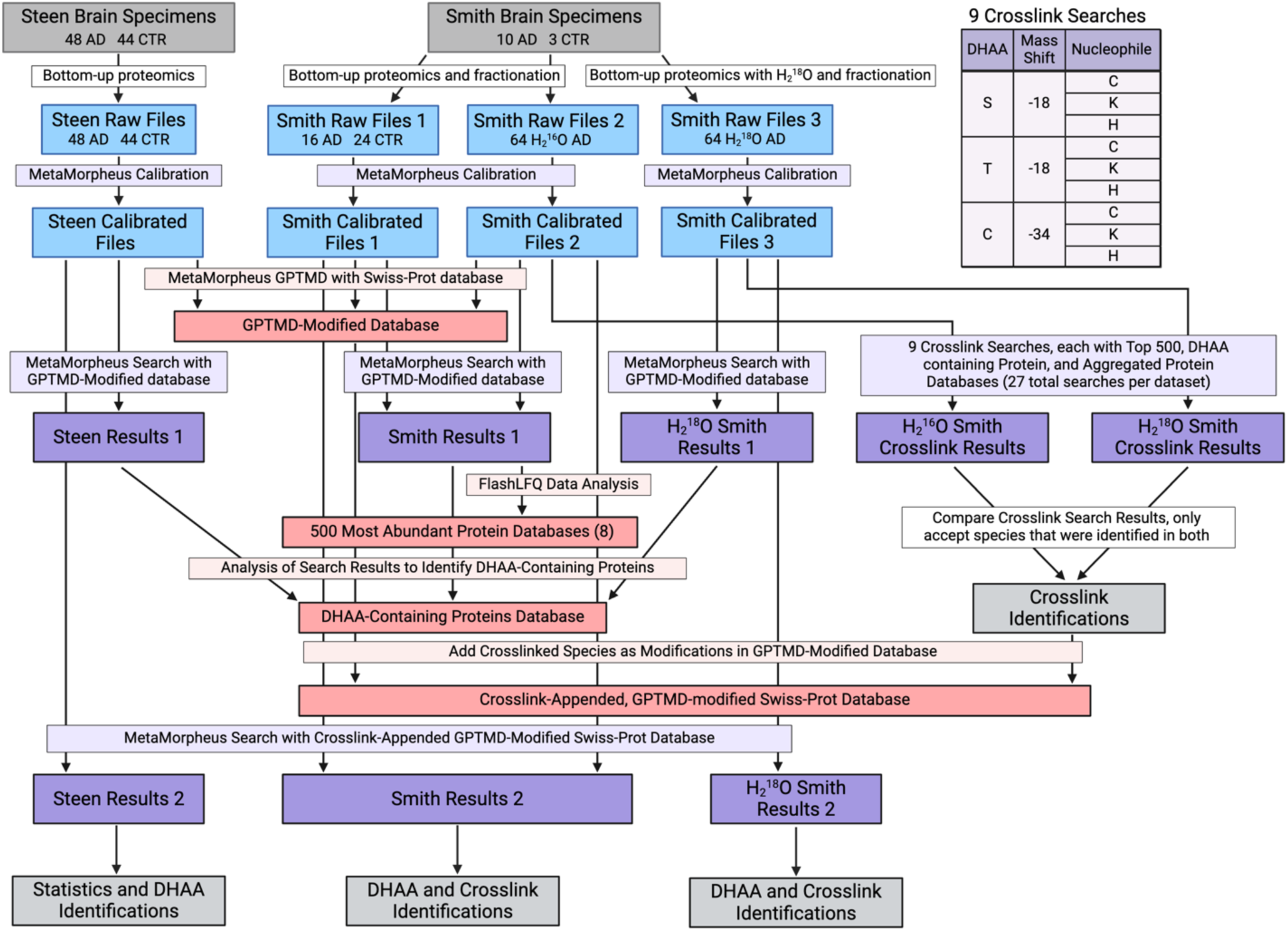
Flowchart for data generation and analysis. Mass spectrometry data is shown in blue. Databases are shown in red. Operations and results generated from MetaMorpheus, our laboratory’s open-source search software, are shown in purple. Specimen inputs and overall result outputs are shown in gray. Briefly, two datasets stemming from the Steen and Smith laboratories were used for this study. Both preparations isolated the sarkosyl-insoluble samples from a number of brain specimens. The Smith dataset utilized fractionation, and thus generated 8 files per brain sample. Smith Raw Files 1 corresponds to BU analysis of 2 AD brains (16 files) and 3 control brains (24 files); Smith Raw Files 2 and 3 correspond to the remaining 8 brains (64 files for each). MetaMorpheus was used to calibrate each raw file. GPTMD, a tool within MetaMorpheus, was used to generate a modified version of the Swiss-Prot human database that included possible PTMs. This database, referred to as the GPTMD-modified database, was used for a standard bottom-up proteomics search. For crosslink searches, nine targeted databases were generated from bottom-up search results, and a tenth targeted database was created consisting of proteins known to aggregate in neurodegenerative disorders. Crosslink searches were performed using these databases. Crosslink identifications were only considered valid if found independently in both the H_2_^16^O and the H_2_^18^O crosslink search results. These validated crosslink identifications were appended to the GPTMD-modified database, and this new database was used to search each data set again. The results reported in this study were obtained from these final searches. Created with BioRender.com.

#### Datasets employed

Two sets of proteomic data were used in this study, one generated within our laboratory (Smith), and one that was previously published by Wesseling et al. and is publicly available (Steen). Our laboratory analyzed the angular gyrus of 10 AD subjects and 3 age-matched controls. Each sample was treated as described above, resulting in 104 raw files (13 samples, 8 fractions each) from standard bottom-up proteomics, and an additional 64 raw files (8 samples, 8 fractions each) from H_2_^18^O treated samples. These files are collectively referred to as the “Smith” data. The datasets generated from H_2_^18^O treated samples (Smith Raw Files 3 in Fig. 2) were used for crosslink identification with the datasets from the H_2_^16^O treated samples prepared from the same brain specimens (Smith Raw Files 2 in Fig. 2). The study by Wesseling *et. al.* analyzed the angular gyrus of 48 AD subjects and 44 age-matched controls, yielding 92 raw files. These files were retrieved from ProteomeXchange and we refer to them collectively as the “Steen” data.

#### Initial bottom-up searches for peptide identifications and PTM discovery

Raw files were analyzed with the open-source search software program MetaMorpheus (https://github.com/smith-chem-wisc/MetaMorpheus). Each file was calibrated using the calibration feature within MetaMorpheus, where the Swiss-Prot human XML (canonical) database was downloaded and used as the reference proteome. Global post-translational modification discovery (GPTMD), a data analysis tool within MetaMorpheus, was used to identify selected PTMs that are not annotated in the reference database and may constitute novel PTM identifications. DHAA conjugates are listed in Table S3 and were added as custom GPTMD modifications allowing them to be searched for on the indicated amino acid residues of all proteins in the database. Modifications identified by GPTMD were appended to the Swiss-Prot reference database, which we refer to as the GPTMD-modified database. The GPTMD-modified database was used to search the Smith and Steen datasets separately. Search results were filtered to a 1% false discovery rate. FlashLFQ (https://github.com/smith-chem-wisc/FlashLFQ) was used to perform quantification of the search results. These results are referred to as Smith and Steen “Results 1” in Fig. 2.

#### Crosslink searches and targeted databases

Crosslink searches were performed using MetaMorpheusXL, a crosslink search tool within MetaMorpheus. Sixteen datasets were used for crosslink searches: 8 for the H_2_^16^O conditions, and 8 for the H_2_^18^O conditions, labeled as Smith Raw Files 2 and 3 in Fig. 2. Crosslink searches often require targeted protein databases to search against in order to keep the search space manageable in size. We generated 10 such targeted databases for this purpose. The first was a set of 22 proteins such as Tau and amyloid-β, that have been reported to aggregate in neurodegenerative disorders. The second was a set of 370 proteins containing DHAAs or their conjugates, generated from the combined Smith and Steen search results described above. The remaining eight targeted protein databases were the 500 most abundant proteins identified in each of the 8 brain specimens, as determined by FlashLFQ.

MetaMorpheusXL allows for custom crosslinkers, which are the combination of the mass of a crosslinker with the sites that are crosslinked. For crosslinks arising from a DHAA from S or T, −18.0105 was used as the crosslinker mass. For crosslinks arising from C, −33.98772 was used as the crosslinker mass. These potential DHAAs were allowed to crosslink with residues with nucleophilic side chains: C, K, and H, which have been reported to react with DHAAs [8,17,27]. This gives a total of 9 crosslink searches (each of the 3 potential DHAAs reacting with each of the 3 possible nucleophilic side chains) to be performed on the 16 fractionated datasets (8 for H_2_^16^O and 8 for H_2_^18^O) with each of the 3 protein databases (aggregated proteins, DHAA-containing proteins, and top 500 proteins by abundance for the respective sample) for a total of (9 × 16 × 3) 432 searches. Note, the searches of H_2_^18^O-labeled Smith data were run with nearly identical parameters, but included a fixed modification for two ^18^O atoms at the C-terminus (+4.008491); the software already takes into account two C-termini on a crosslinked pair of peptides resulting in the ∼8 Da mass difference compared to the H_2_^16^O search results.

Results from the 432 crosslink searches were pooled to create four files: Intralinks Standard, Interlinks Standard, Intralinks H_2_^18^O-treated, and Interlinks H_2_^18^O-treated, where Intralinks refer to crosslinks between the same protein species (e.g. Tau S262-C291) and Interlinks refer to crosslinks between different protein species (e.g. GAPDH T246-VDAC2 H284). These files are referred to as “Smith Crosslink Results” in Fig. 2. Results were compared manually to determine crosslinks that were identified in both standard and H_2_^18^O-treated datasets. Fragmentation was analyzed in the MetaMorpheus visualization software. MS1 verification was performed manually in QualBrowser (Thermo Fisher Scientific) and compared to theoretical isotope distributions generated by Protein Prospector (UCSF, http://prospector.ucsf.edu).

#### Final bottom-up proteomics search with crosslink-appended,GPTMD-modified Swiss-Prot database

The GPTMD-modified database used for previous MetaMorpheus searches was duplicated and appended with crosslinks discovered from MetaMorpheusXL searches. These crosslinks were added as custom modifications at the residue sites where they were observed, as listed in Table S3. This database is referred to as the “Crosslink-Appended, GPTMD-modified Swiss-Prot Database” in Fig. 2. Using this database, the Smith and Steen data were again searched separately, and label-free quantification was performed separately in FlashLFQ. An analogous database was created for the H_2_^18^O-treated datasets by specifying that custom crosslink modifications contain an ^18^O-labeled C-terminus (+4.00849). Calibration and search tasks were edited to contain a fixed modification for ^18^O-labeled C-termini (+4.00849) and the H_2_^18^O-treated datasets were searched together. Lists of dehydroamino acids and their derivatives, including crosslinks, were generated from these three final result files, called “Results 2” in Fig. 2 (Data S1).

#### Searches of sarkosyl-soluble data

The non-aggregate-enriched sarkosyl-soluble data was searched two ways. First, the samples were calibrated and analyzed with GPTMD and the Swiss-Prot standard database as before. Searches were conducted under the same parameters as the sarkosyl-insoluble datasets. Second, the Crosslink-Appended, GPTMD-modified Swiss-Prot Database was used to determine if crosslinks were present in these samples. DHAA identifications from these searches are available in Data S2.

#### Statistics

A two-tailed t-test was employed to evaluate confidence for the differential abundance data presented in Fig. 4A. The Steen data, obtained from 48 AD samples and 44 control samples, were used for this analysis. Error bars for the occupancy data in Fig. S11 and Table S4 were also calculated using the Steen data, using standard error of the mean and its propagation.

#### Spike-in experiments

Data obtained from spike-in experiments were searched using the Crosslink-Appended, GPTMD-modified Swiss-Prot database. Label free quantification was used to estimate the relative abundance of each crosslinked species. Extracted ion chromatograms were generated for each crosslinked species, and MS1 spectra were averaged over the period of elution. Intensities of the crosslinked species in the presence or absence of spiked-in standards were determined.

## Results

### Evidence for dehydroamino acids in the sarkosyl-insoluble proteins from Alzheimer’s disease

We utilized a high-quality mass spectrometry proteomics dataset from Wesseling *et al.,* produced in a study directed towards identifying PTMs of the Tau protein in patients with and without AD [23]. The authors prepared the sarkosyl-insoluble proteins from brain specimens, effectively enriching for the protein aggregates in which hypothesized DHAAs might be most abundant. They analyzed both sarkosyl-soluble and sarkosyl-insoluble fractions. They also collected high resolution tandem MS data, which is invaluable for deep characterization of PTMs in complex samples. In total, this study of 49 AD and 44 control brain specimens identified 95 unique PTMs of Tau.

Our initial searches of these data using the MetaMorpheus search engine to look for DHAAs and their conjugates revealed many instances in these brain specimens. This is, to our knowledge, the first evidence for DHAAs or their conjugates in AD. To follow up on these results from public data, the angular gyrus sections of ten AD brain specimens and three age-matched controls were obtained from the Wisconsin Alzheimer’s Disease Research Center (WADRC). Sarkosyl-soluble and insoluble fractions were prepared as described in Methods. After tryptic digestion to generate peptides, high-pH offline fractionation was employed to yield 8 fractions per insoluble biological sample, thereby increasing proteomic coverage depth.

MetaMorpheus search results of the combined Wesseling *et al.* and our own datasets revealed many DHAAs and DHAA conjugates. We searched 48 AD and 44 control sarkosyl-insoluble samples and 28 AD and 28 control sarkosyl-soluble samples from the Wesseling *et al.* datasets. We further searched 10 AD and 3 controls for both sarkosyl-insoluble and soluble material generated by our laboratory, with the insoluble material being fractionated for deeper coverage. An order of magnitude more DHAAs were identified in the sarkosyl-insoluble material than in the sarkosyl-soluble material, as shown in Table 1. A similar order-of-magnitude difference is seen if only the Wesseling *et al.* data are employed for the comparison, although the absolute numbers of identifications are lower (Table S5). We searched for six different forms at each eliminylation site (Fig. 1): DHAA, GSH-, HC-, dithiothreitol (DTT)-, and tris(2-carboxyethyl)phosphine (TCEP)-modified DHAAs, as well as DHAA-XLs (see Methods). Other DHAA reaction products may also be present in the samples, but our study focused on these likely conjugates (Text S1). We required that at least one conjugate species (GSH-, HC-, DTT-, TCEP-, or DHAA-XL) be observed in order to confidently identify a site where DHAAs are generated, as the direct observation of DHAA itself with a −18/-34 Da mass shift of H_2_O/H_2_S loss may be generated artifactually [13]. We did not also require that the DHAA itself be directly observed at that site, as the mass shift signature of the DHAA conjugate is a more reliable indicator. We refer to these identifications as eliminylation sites, and they are reported in Data S1 and S2 for sarkosyl-insoluble and -soluble datasets.

**Table 1:** Numbers of eliminylation sites and proteins containing DHAA conjugates.

|  | Sarkosyl-Insoluble |  | Sarkosyl-Soluble |  |
| --- | --- | --- | --- | --- |
|  | Sites | Proteins | Sites | Proteins |
| Any Identification | 681 | 266 | 37 | 28 |
| >50% of Specimens* | 412 | 184 | 25 | 17 |
| >80% of Specimens | 185 | 84 | 13 | 13 |
| 100% of Specimens | 42 | 22 | 1 | 1 |

\*For consideration in this manuscript regarding the number of DHAA modifications observed, we somewhat arbitrarily set the threshold for considering a DHAA to have been widely identified to be that it was observed in at least 50% of all specimens analyzed. This corresponds to the ">50% of Specimens" entry above, i.e. 412 sites in 184 proteins.

Table 2 illustrates that several eliminylated proteins are either involved directly in AD pathogenesis or related to a pathway that is dysregulated in AD [28–32]. Most notably, several aggregation-prone proteins in AD including Tau (MAPT), CRMP2 (DPYSL2), and α-synuclein (SNCA) contain sites of eliminylation. We note that amyloid-ß was found to contain two DHAAs, but no conjugates were identified to permit their validation (Data S1).

**Table 2:**
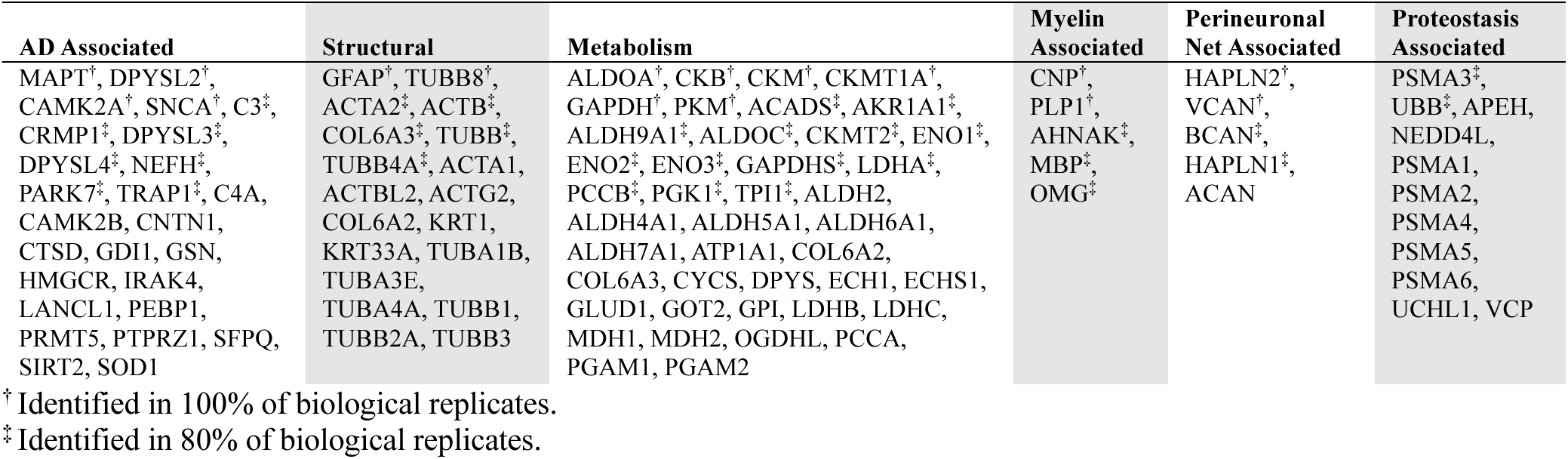
Dehydroamino acid modified proteins identified in 50% or more of specimens.

### Dehydroamino acids produce crosslinks in Tau

Dehydroamino acids can react with nucleophilic side chains of cysteine, lysine, and histidine to yield protein-protein crosslinking [27]. This reaction is analogous to that which occurs for DHAAs with small molecule biological nucleophiles such as glutathione and homocysteine, where the thiol readily reacts with the electrophilic alkene in DHAAs. We looked for evidence of DHAA-mediated protein-protein crosslinking using MetaMorpheusXL, a specialized software tool within MetaMorpheus that was developed specifically for the identification of crosslinked peptides [33]. Our own data, plus that from Wesseling *et al.*, was searched initially against protein databases comprised of the major aggregating proteins in AD. The choice of a small, targeted protein database for crosslink searches is essential, as the search space increases rapidly with increasing numbers of protein entries, increasing both the likelihood of false positive identifications and run time. MetamorpheusXL searches yielded evidence for two dehydroamino acid-mediated crosslinks in the Tau protein: S262-C291 and S400-C291. Notably, S262 and S400 were also identified as eliminylation sites in the MetaMorpheus search results presented above, consistent with the premise that the observed crosslinks had formed from DHAA precursors. The S262-C291 and S400-C291 crosslinks were identified with peptide precursor mass errors of −0.29 and −1.4 ppm, respectively. An example MS2 fragment spectrum for the S400-C291 crosslink is presented in Fig. 3A. This spectrum provides strong evidence for a crosslink between S400 and C291: (i) thirteen y-ions and three b-ions are observed to match theoretical fragments from the crosslinked species, (ii) three y-ions and one b-ion include the crosslink in the fragment, meaning the entire mass of the “other” peptide is observed as a modification mass on the requisite fragment, and (iii) detected fragments localize the crosslink to S400 and C291, as a linkage anywhere else in the respective peptides would not give the observed fragmentation pattern.

**Fig. 3:**
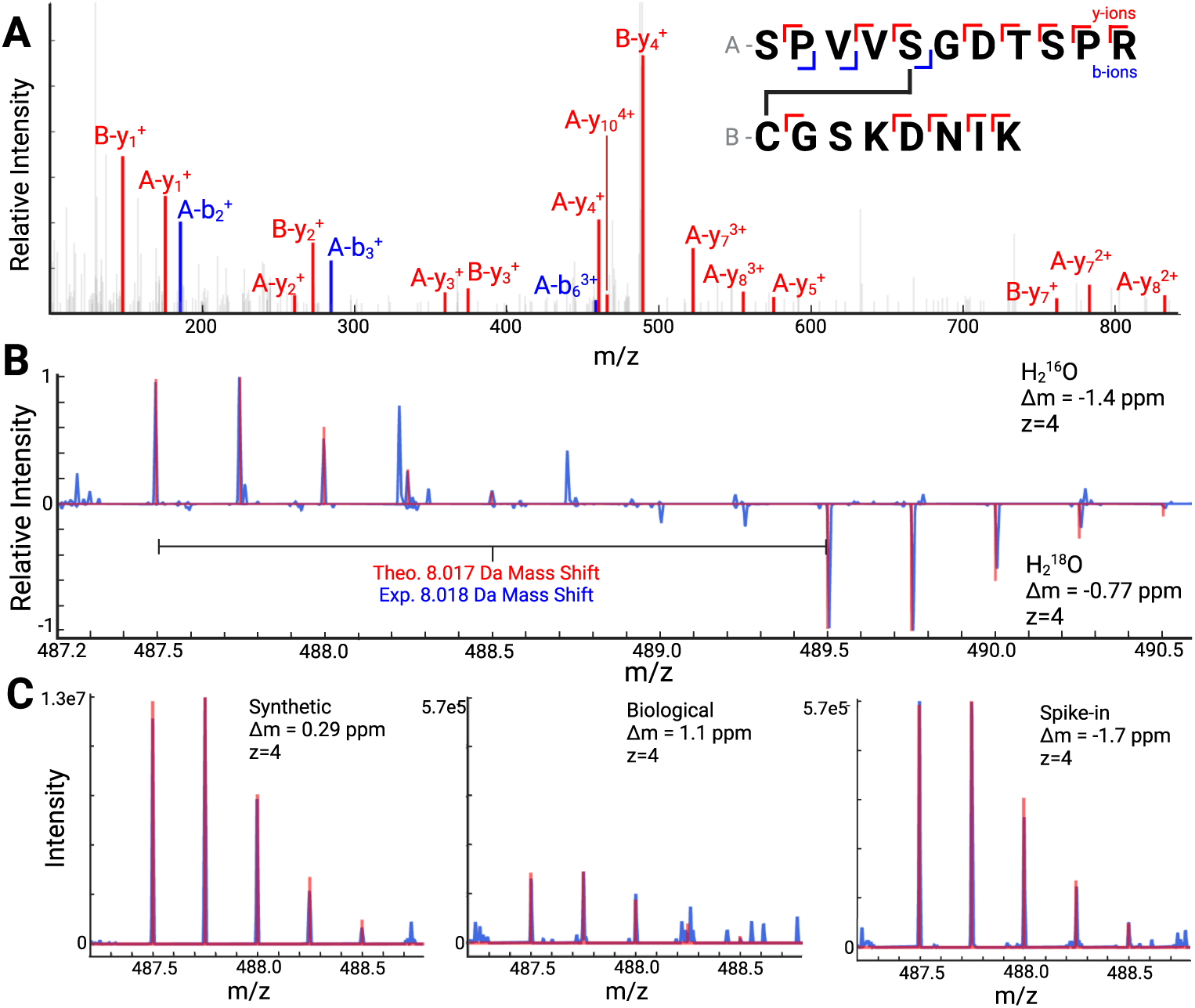
Evidence for a DHAA-mediated crosslink between residues S400-C291 in Tau. **A**, The MS2 fragmentation spectrum for the precursor mass corresponding to the crosslinked species. It shows thirteen y-ions and three b-ions, including four especially important fragments (A-y_7_, A-y_8_, A-y_10_ and A-b_6_) that include the crosslink. Fragmentation throughout the peptide is sufficient to accurately assign the crosslink to residues S400 and C291. **B**, A mirror plot is shown with MS1 data derived from H_2_^16^O samples on top (usual preparation) and H_2_^18^O samples on the bottom (heavy isotope preparation). Raw data are shown in blue, and the theoretical isotope distributions are overlaid in red. The close match for each isotope peak compared to its theoretical value, for both mass and relative intensity, supports that the crosslink peptide has been identified correctly. H_2_^18^O labeling validates the presence of a crosslink between two peptides containing residues S400 and C291 in Tau. **C**, MS1 spectra for the synthetic standard, the biological sample, and the spike-in are shown with raw data in blue and theoretical isotope distributions overlaid in red. Created with BioRender.com and Plotly [34]. Evidence for the S262-C291 crosslink, as well as other crosslinks, can be found in Fig. S1-10.

### Validation of the Tau crosslink using H_2_^18^O labeling

To confirm further that the crosslinked peptides observed were correctly identified, we employed a previously established C-terminus isotopic labeling strategy [24]. Briefly, whereas standard linear tryptic peptides contain only one C-terminus with two C-terminal oxygen atoms, crosslinked tryptic peptides will contain two C-termini, and thus four C-terminal oxygen atoms. Replacing a standard tryptic digestion buffer with buffered H_2_^18^O leads each C-terminal oxygen to be replaced with isotopically distinct ^18^O, a mass shift of two daltons compared to the normal ^16^O isotope. The observed mass of crosslinked peptides will therefore be shifted by +8 Da, whereas standard peptides will be shifted by +4 Da, clearly indicating whether the observed species is crosslinked or not. The sarkosyl-insoluble portions of eight AD brain specimens were prepared as described in Methods and digested with trypsin in H_2_^18^O. The expected mass shift of +8 Da was observed for the database search identifications of both the S262-C291 and the S400-C291 crosslinks, confirming their identities as crosslinked species. The +8 Da mass shift is shown in the MS1 spectra in Fig. 3B comparing H_2_^16^O and H_2_^18^O preparations for the S400-C291 crosslink.

### Synthetic crosslinked peptide standards corroborate dehydroamino acid-mediated crosslinks in Tau protein

To unequivocally confirm these Tau crosslink identifications, crosslinked peptide standards were synthesized and then analyzed by mass spectrometry. The constituent tryptic peptides for each crosslink were purchased with a phosphorylation modification at the site of eliminylation. The phosphoserine was eliminylated using Ba(OH)_2_ to generate the dehydroamino acid [26]. The eliminylated peptide was incubated with the corresponding cysteine-containing peptide to generate the crosslinked species. MS analysis of these synthesized crosslinked peptide standards supports the identification of S262-C291 and S400-C291 DHAA-XLs in Tau, as the observed intact masses and isotopic distributions, as well as the observed fragmentation patterns in MS2 spectra all match those of the identified crosslinked species in the AD brain specimens.

Having determined that the MS data for our synthetic standards matched the native species, we further verified the identity of the biological species through a spike-in experiment. The synthetic crosslinked peptides were added to a prepared brain sample where both S262-C291 and S400-C291 had been observed previously by mass spectrometry. The data from the spike-in sample displayed an increase in all three metrics of analysis for each of the two crosslinked species: (i) the intensity of the extracted ion chromatogram, (ii) the intensity of each peak in the intact mass isotopic distribution of the MS1 spectrum, and (iii) the intensity of each fragment observed by MS2 analysis. The MS1 spectra corresponding to the S400-C291 crosslink are presented in Fig. 3C. This spike-in experiment provides irrefutable evidence of the existence of two DHAA-XLs in Tau, between residues S262-C291 and S400-C291.

### Discovery of dehydroamino acid-mediated crosslinks in other proteins

In addition to the observations of crosslinks in the Tau protein described above, crosslinks were also observed in other proteins present in the AD and control brain specimens. These crosslink identifications were based on three criteria. First, crosslinks must be validated by ^18^O labeling, meaning the crosslinked species are found in the standard bottom-up samples with mass M, and in the ^18^O treated samples with mass M+8, within an acceptable retention time window. Only crosslinks identified in both standard preparations and with the +8-Da shift in the H_2_^18^O-datasets were accepted. Second, masses obtained from MS1 spectra must match the theoretical mass within an acceptable range. Lastly, the DHAA-XLs (both standard and H_2_^18^O forms) must be reported below a 1% FDR in a search against the full proteome. To determine if DHAA-XLs met this criterion, crosslinks were entered as modifications in the MetaMorpheus Global Posttranslational Modification Discovery (GPTMD) tool [35,36], and searched using a standard bottom-up proteomics search. See Methods for more information regarding these searches and filtering criteria.

Eleven total DHAA-XLs, including two previously detailed in Tau, met these criteria (Table 3). Mass spectrometry data support these findings (Fig. 3, Fig. S1-10). Several crosslinks, including those identified within pathogenic proteins such as amyloid-β, were excluded from Table 3 as they did not satisfy one or more of the predefined criteria. Interested readers may refer to Table S6 for a comprehensive list of these additional crosslinks.

**Table 3:**
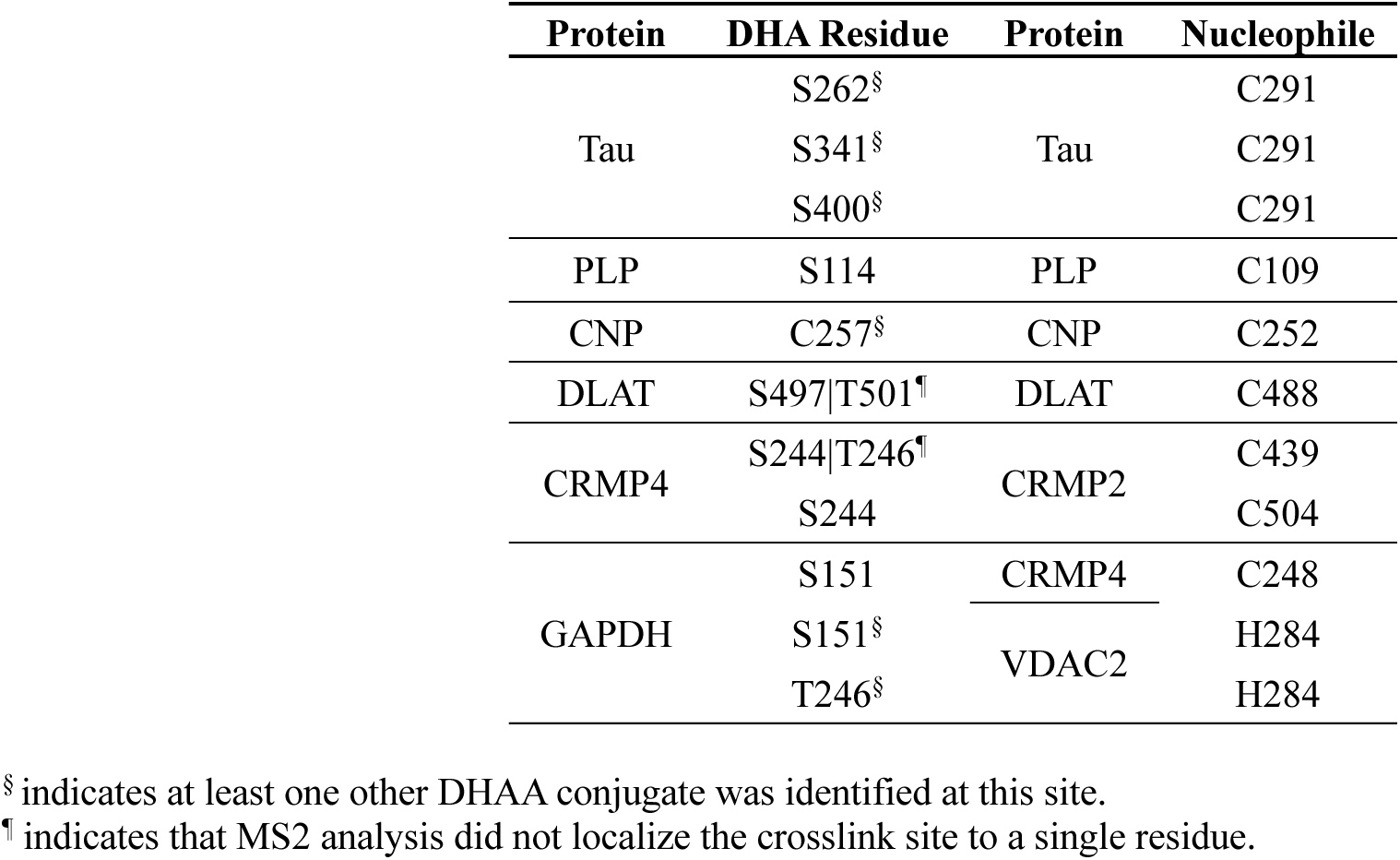
Dehydroamino acid-mediated crosslinks. The Tau, PLP, CNP and DLAT crosslinks may be either intra- or intermolecular, but we highlight that it is possible that these crosslinks are intermolecular, as evidenced by crosslinks between CRMP4, CRMP2, GAPDH, and VDAC2, which are all necessarily intermolecular.

| Protein | DHA Residue | Protein | Nucleophile |
| --- | --- | --- | --- |
| Tau | S262<sup>§</sup> | Tau | C291 |
|  | S341<sup>§</sup> |  | C291 |
|  | S400<sup>§</sup> |  | C291 |
| PLP | S114 | PLP | C109 |
| CNP | C257<sup>§</sup> | CNP | C252 |
| DLAT | S497\|T501<sup>¶</sup> | DLAT | C488 |
| CRMP4 | S244\|T246<sup>¶</sup> | CRMP2 | C439 |
|  | S244 |  | C504 |
| GAPDH | S151 | CRMP4 | C248 |
|  | S151<sup>§</sup> | VDAC2 | H284 |
|  | T246<sup>§</sup> |  | H284 |

<sup>§</sup> indicates at least one other DHAA conjugate was identified at this site.
<sup>¶</sup> indicates that MS2 analysis did not localize the crosslink site to a single residue.

### Quantitative comparisons involving DHAA modifications

The three most heavily DHAA-modified proteins observed in this study were GAPDH, CRMP2, and Tau (Data S1). GAPDH is one of the very small number of human proteins previously reported to undergo DHAA modification [12]; while CRMP2, like Tau, is a hyperphosphorylated, NFT-forming microtubule associated protein [3]. Quantitative analysis of the relative abundance in AD compared to control specimens is shown in the volcano plot in Fig. 4A for all phosphorylation, DHAA, and DHAA-conjugate modifications. The modifications of Tau stand out in this plot, as they are most clearly present at higher levels in AD than control brains, consistent with the premise that they have an important role in the disease state. It is noteworthy that the observed fold changes and associated p-values for the Tau DHAA modifications are comparable to those for Tau phosphorylation events, which are widely recognized as important to the aggregation of Tau, and thus to AD pathology [5]. We note that phosphorylation of Tau T231 was the most confident differentially abundant PTM observed, followed by the Tau crosslink between S262-C291 (Fig. 4A). Phosphorylation at T231 has been shown to have profound structural effects on Tau and has been proposed as a cerebrospinal fluid biomarker for AD [37,38], suggesting that the crosslink between S262-C291 may have similar importance. The other two Tau crosslinks, S400-C291 and S341-C291, were also found to be far more abundant in AD aggregates than in controls. Table S7 presents the fold-changes and p-values for each DHAA site identified. The site occupancies of DHAAs and DHAA conjugates at each eliminylation site are estimated in Fig. S11 and Table S4. The locations of both the Tau eliminylation sites and the Tau phosphorylation sites are shown in Fig. 4B with the majority of sites being shared. This observation is intriguing, as phosphorylation is a likely precursor to DHAA formation [7].

**Fig. 4:**
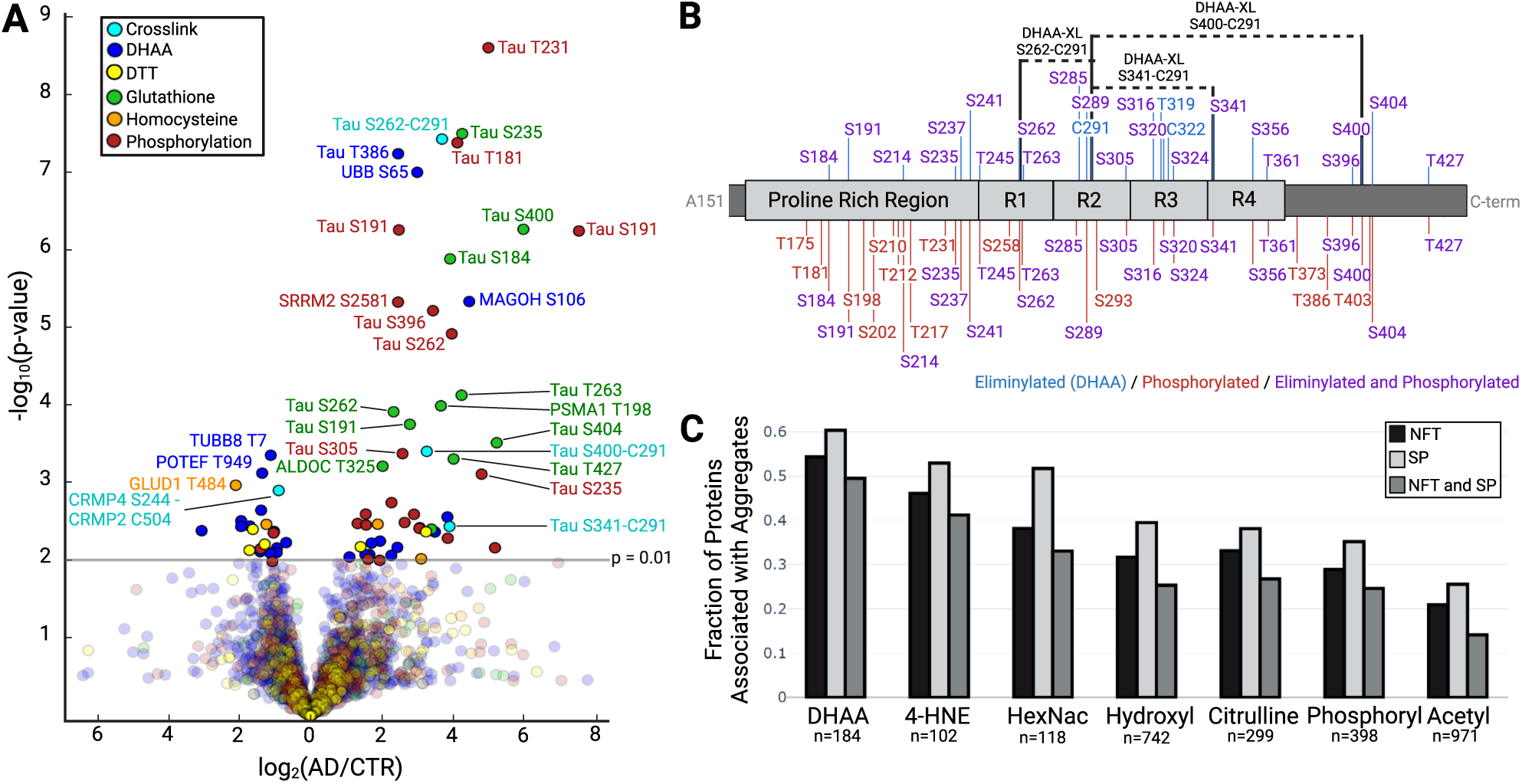
**A**, A volcano plot showing the differences between AD and control brain specimens for DHAAs, their conjugates, and phosphorylation. The fold changes and associated p-values for DHAA-related modifications in Tau are comparable to those for well-documented phosphorylation events. Most notably, Tau crosslinks are much more prevalent in AD than age matched controls (S262-C291–12.8-fold higher, p=3.4e-8; S400-C291–9.55-fold higher, p=3.7e-4; and S341-C291–14.8-fold higher, p=3.5e-3). **B**, DHAA sites identified by a conjugate in 50% or more of brain specimens are plotted (blue lines, top) along with the sites of phosphorylation (red lines, bottom). Three of these sites, S262, S341, and S400, form inter- or intramolecular crosslinks with Tau C291, represented by the dotted black lines. **C**, Fraction of proteins involved in NFTs and SPs, sorted by their observed PTMs (Table S8). Created with BioRender.com and Plotly [34].

The increased levels of Tau DHAAs in AD brain specimens compared to age-matched controls suggests a possible role for dehydroamino acids in the pathogenesis of AD. We sought to determine how the occurrence of this modification compares to that of other PTMs present in brain protein aggregates. It has been shown that NFTs and SPs, while composed primarily of Tau and amyloid-ß respectively, are also commonly associated with numerous secondary proteins. Ferreira *et al.* identified these secondary proteins in a bioinformatic analysis, associating 627 proteins with NFTs and 857 proteins with SPs [39]. We identified 418 of these 627 NFT-associated proteins and 555 of the 857 SP-associated proteins in our study. We sorted all identified proteins into subset lists by observed PTMs (e.g. all phosphorylated proteins comprise the “Phosphorylation” list). Each list was compared to the aggregate-associated proteins assigned by Ferreira *et al.*, and the fraction belonging to those NFT and SP sets was determined for each PTM type. The results are presented in Fig. 4C. A larger proportion of the observed DHAA-modified proteins are associated with AD-specific protein aggregation than any other individual PTM, providing additional support for the hypothesis that DHAAs are related to protein aggregation in AD.

## Discussion

In this study we have shown that dehydroamino acids, their conjugates, and the crosslinks that they form comprise a rich and previously unknown source of proteomic complexity in Alzheimer’s disease brain specimens. The inherent nature of DHAAs, either through their potential to change protein secondary structure or their ability to form protein-protein crosslinks, provide an obvious potential connection to protein aggregation. A possible role for DHAAs in the protein aggregation processes of AD is supported by previous studies implicating them in the aggregation of the αB-crystallin protein in the human eye lens. Support for this hypothesis is also provided by studies showing that overexpression of the DHAA-modifying enzyme LanCL1 attenuates effects of aging, oxidative stress, and neurodegenerative disease in murine brain models [20–22]. We note that several current therapeutic candidates may affect DHAAs (Text S2). The origin of the observed DHAA modifications remains unknown, although both oxidative damage and enzymatic routes have been suggested [17–19]. We are currently searching for a human enzyme capable of catalyzing eliminylation to generate DHAAs.

The results presented above provide a strong connection between DHAA prevalence and protein aggregation, a hallmark of Alzheimer’s disease. We confidently identified 412 eliminylation sites in 184 proteins from the protein aggregate-enriched sarkosyl-insoluble material from AD-brain specimens. In contrast, only 25 eliminylation sites in 17 proteins were identified in the sarkosyl-soluble material, showing that DHAAs are highly enriched in protein aggregates (Table 1). The quantitative differences in DHAA levels between AD and controls indicate a potential role for these modifications in AD pathology. The DHAA-containing proteins are associated with key cellular functions, including metabolism, myelin structure and maintenance, perineuronal net construction, and proteostasis, all of which are dysregulated in AD [28–32]. The DHAA-modified proteins were more strongly associated with NFTs and SPs than were proteins with any other type of PTM. Many of the same DHAAs and crosslinks observed in the AD samples were also detected in sarkosyl-insoluble samples from age-matched controls, albeit at a lower abundance. We hypothesize that this may reflect a lower level of DHAA modification and/or protein aggregation preceding symptomatic onset in AD [40].

We identified eleven dehydroamino acid-mediated crosslinks in eight proteins. These proteins include Tau, CRMP2, and CRMP4, which are major constituents of NFTs in AD [3]. We also found these crosslinks in the myelin proteins PLP1 and CNP; while these proteins have not been reported to aggregate, a possible connection is suggested by a recent study showing that amyloid-associated microglial cells meant to clear aggregated proteins respond to myelin damage [41]. GAPDH, which is heavily modified with DHAAs and which we found to form crosslinks with CRMP4 and VDAC2, is well documented as a co-aggregating species in AD [42]. We do not yet know if the observed DHAA-XLs in Tau, PLP1, CNP and DLAT occur intra- or intermolecularly. Further work will seek to address this question, but we note that at least some of the DHAA crosslinks identified here are necessarily intermolecular, as they link two different proteins (e.g. CRMP2 and CRMP4).

## Conclusions

This study provides evidence that dehydroamino acids are a highly prevalent modification in protein aggregates formed in Alzheimer’s disease. The demonstrated correlation of DHAAs with protein aggregates, as well as the significant differential abundance of DHAA-related species in AD, suggests that this relatively obscure modification could be playing an important role in protein aggregation. While this study focused on AD brain specimens, the important role that has been shown for LanCL1 in other neurodegenerative disorders, such as Parkinson’s disease [22] and Amyotropic Lateral Sclerosis [21], suggests that DHAA-related modifications may be prevalent in those systems as well. We hope that this discovery of a potential new basis of protein aggregation will spur investigations into DHAAs in neurodegenerative disorders that eventually ameliorate the consequences of these diseases.

## Supporting information

Text S1-2, Figures S1-S11, Tables S1-2, S5-6, and Schemes S1-2

Data S1 - Insoluble Bioreps

Data S2 - Soluble Bioreps

Table S3 - Modifications

Table S4 - Occupancy

Table S7 - Differential Abundance

Table S8 - PTMs v. NFT and SP

## List of abbreviations

ACN: Acetonitrile
AD: Alzheimer’s disease
DHAA-XL: Dehydroamino acid-mediated crosslink
DHAA: Dehydroamino acid
DTT: Dithiothreitol
GPTMD: Global posttranslational modification discovery
GSH-DHAAs: Glutathionylated dehydroamino acids
GSH: Glutathione
HC: Homocysteine
HPLC: High performance liquid chromatography
IRB: Institutional review board
MPTP: 1-methyl-4-phenyl-1,2,3,6-tetrahydropyridine
MS: Mass spectrometry
NFT: Neurofibrillary tangle
PTM: Posttranslational modification
SP: Senile plaque
TCEP: tris(2-carboxyethyl)phosphine
WADRC: Wisconsin Alzheimer’s Disease Research Center
WBDP: Wisconsin Brain Donor Program

## Declarations

### Ethics approval and consent to participate

Since this study used only decedent tissue samples and the scope was consistent with WBDP protocols, this study was determined to meet the standards of minimal risk and was exempted from local IRB oversight after their review.

### Consent for publication

Not applicable.

### Availability of data and materials

Mass spectrometry data has been deposited to the ProteomeXchange Consortium via the PRIDE partner repository with the data set identifier PXD053814. For reviewer access to these data, login to the PRIDE repository using project accession “PXD053814” with token “l8z22bvmnkcy” or login with and password “O7Twr1Zrxl8c”.

### Author’s contributions

S.W.M, B.L.F., and L.M.S. conceived the idea for the project; All authors contributed to securing funding for this work; S.W.M., B.L.F., M.S., and L.M.S. designed methodology; S.W.M. and M.S. performed experiments; S.W.M., B.L.F., and M.R.S. performed data analysis; S.W.M., B.L.F., and L.M.S. wrote the paper with input from all authors.

## Acknowledgements

We thank James W. Bruce, Mark Horswill, and Masaki Nishikiori for ultracentrifuge access and guidance. We thank Nicholas E. Bollis and Alexander J. Solivais for their advice in data analysis. We would like to express our appreciation for the very high quality and comprehensive proteomic data made publicly available by the Judith Steen laboratory (Wesseling et al.), without which this work would have been substantially more limited. We thank the Wisconsin Alzheimer’s Disease Research Center for providing the brain tissue samples employed in this study.

