## Supplementary material for "Discovery of dehydroamino acids and their crosslinks in Tau and other aggregating proteins of Alzheimer’s disease": Text S1-2, Figures S1-S11, Tables S1-2, S5-6, and Schemes S1-2

**This Supplementary Material 1 file includes:**

Text S1 and S2

Figures S1-S11

Tables S1, S2, S5, and S6

Schemes S1 and S2

**Other Supplementary Materials for this manuscript include the following:**

Data S1 and S2 can be found in Supplementary Materials 2 and 3, respectively.

Tables S3, S4, S7, S8 can be found in Supplementary Materials 4, 5, 6, and 7, respectively.

**Text S1:**

We chose to search for DHAAs and five DHAA conjugates: GSH-DHAA, HC-DHAA, DTT-DHAA, TCEP-DHAA, and DHAA-XLs. The DHAA-XLs were searched for separately in crosslink searches and then added as modifications on particular residues, as described in Methods. DTT and TCEP are both nucleophiles added as reducing agents (DTT in samples prepared by Wesseling *et. al.* [23] and TCEP in samples prepared by our laboratory). We chose to include two biological small molecules, glutathione and homocysteine, as possible nucleophiles that may react with DHAAs, though several other biological nucleophiles exist in the human brain. We sought to limit our search space by limiting the number of conjugates, ultimately resulting in a better false discovery rate. We prioritized the most abundant and most nucleophilic molecules. Glutathione is the most abundant small molecule thiol in human cells [43]. It was also recently reported that the human protein LanCL catalyzes the addition of glutathione to DHAAs [15], therefore making it important to include in our search for DHAA conjugates. Homocysteine was included because it is another high abundance thiol-based nucleophile. There is also evidence that homocysteine accumulates in the brain of patients with AD and may contribute to the pathogenesis [44]. There are several amine-based nucleophiles, including amino acids and biogenic amines such as dopamine, norepinephrine, epinephrine, histamine, and serotonin [45]. While these are nucleophilic and could react with DHAAs, they are less likely to react than glutathione or homocysteine due to the lower nucleophilicity of amines compared to thiols. The reactions of these other nucleophiles with DHAAs lies beyond the scope of this study but may be an area for further investigation.

**Text S2:**

While there are no therapeutics that target DHAAs directly, several therapeutic options are currently being tested for efficacy in AD that may inadvertently affect DHAA formation or its downstream reactivity. When DHAAs are enzymatically generated in microorganism proteins, they arise from phosphorylated precursors [7]. Therefore, preventing the phosphorylation event at eliminylation sites may prevent DHAA formation. Several kinase inhibitors are currently under investigation for their efficacy in treating AD. These kinase inhibitors may target GSK [46], Cdk5 [47], JNK3 [48], p38 MAPK [48], and MEK1/2 [49], all of which are known to contribute to AD pathogenesis and phosphorylate Tau, where we find numerous eliminylation sites. Therefore, by preventing these kinases from acting on Tau, they may prevent DHAA formation. We note that a direct inhibitor cannot yet be designed for a hypothetical DHAA-generating enzyme, as there is no such enzyme known in humans to catalyze this reaction (though some are suspected) [18,19].

Another therapeutic option of note is glutathione (or similar derivatives), which are common antioxidants [50]. These supplements are meant to treat the detrimental oxidative stress observed in AD. While glutathione has many roles in the cell, the human enzyme LanCL is known to add glutathione to DHAAs [15]. Thus, increasing glutathione concentrations by administering a related supplement may affect this pathway by increasing glutathionylation of DHAAs by LanCL, and thereby prevent downstream crosslinking or other potentially detrimental DHAA conjugations.

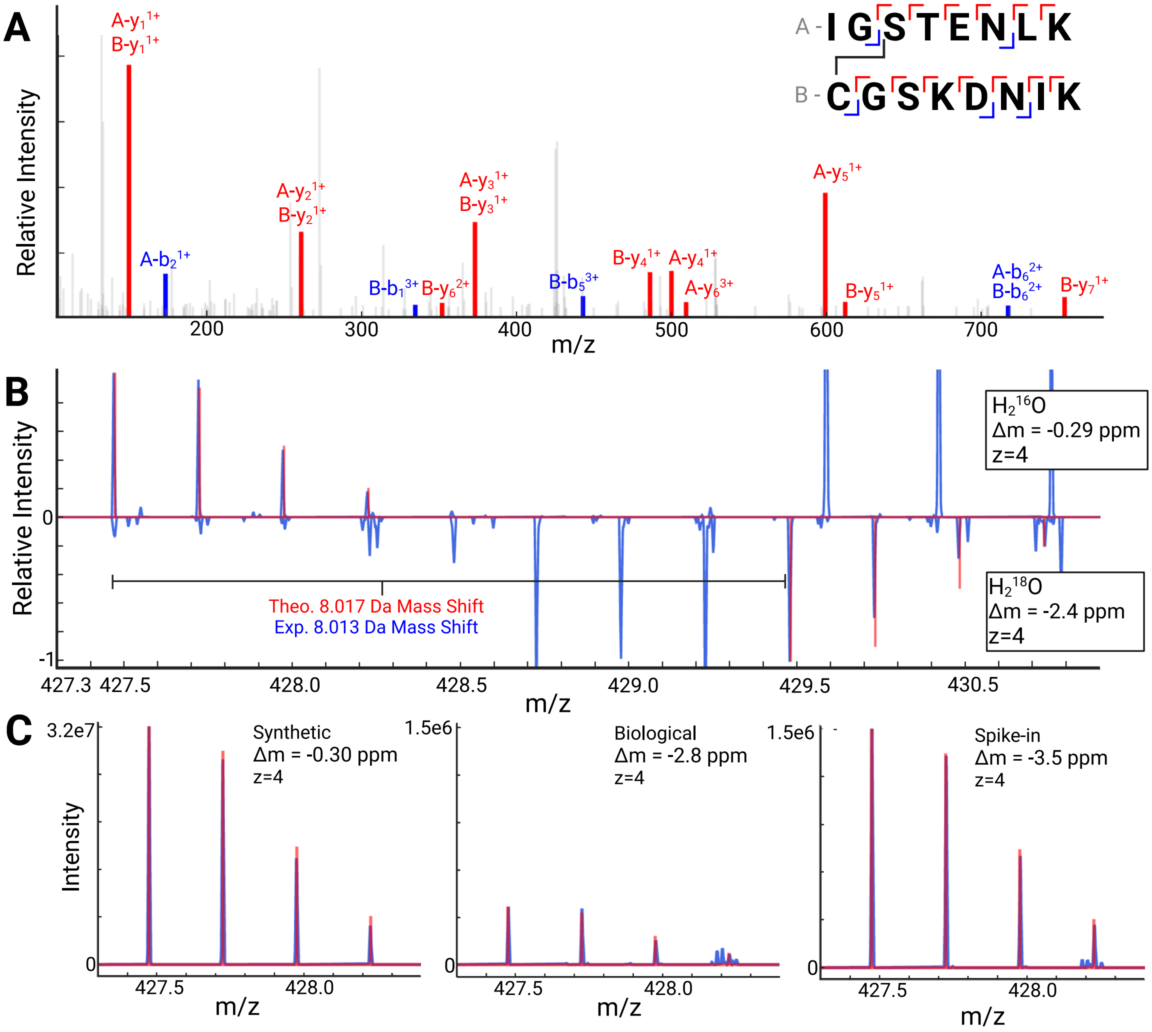


**Fig. S1: Evidence for a DHAA-mediated crosslink between residues S262-C291 in Tau**. **A**, The MS2 fragmentation spectrum for the precursor mass corresponding to the crosslinked species. It shows thirteen y-ions and five b-ions, including five especially important fragments (A-$y_6$, A-$b_6$, B-$b_1$, B-$b_5$, and B-$b_6$) that include the crosslink. Fragmentation throughout the peptide is sufficient to accurately assign the crosslink to residues S262 and C291. **B**, A mirror plot is shown with MS1 data derived from $H_2^{16}O$ samples on top (usual preparation) and $H_2^{18}O$ samples on the bottom (heavy isotope preparation). Raw data are shown in blue, and the theoretical isotope distributions are overlaid in red. The close match for each isotope peak compared to its theoretical value, for both mass and relative intensity, supports that the crosslink peptide has been identified correctly. Additional peaks in the $H_2{}^{16}O$ spectrum are from coeluting peptide species. Additional peaks in the $H_2{}^{18}O$ spectrum are in part from coeluting species, but also from incomplete isotopic labeling of the C-terminal oxygens. $H_2{}^{18}O$ labeling validates the presence of a crosslink between two peptides containing residues S262 and C291 in Tau. C.) MS1 spectra for the synthetic standard, the biological sample, and the spike-in are shown with raw data in blue and theoretical isotope distributions overlaid in red. These data provide unequivocal evidence for a crosslink between S262 and C291 in Tau. Created with BioRender.com and Plotly.[34]

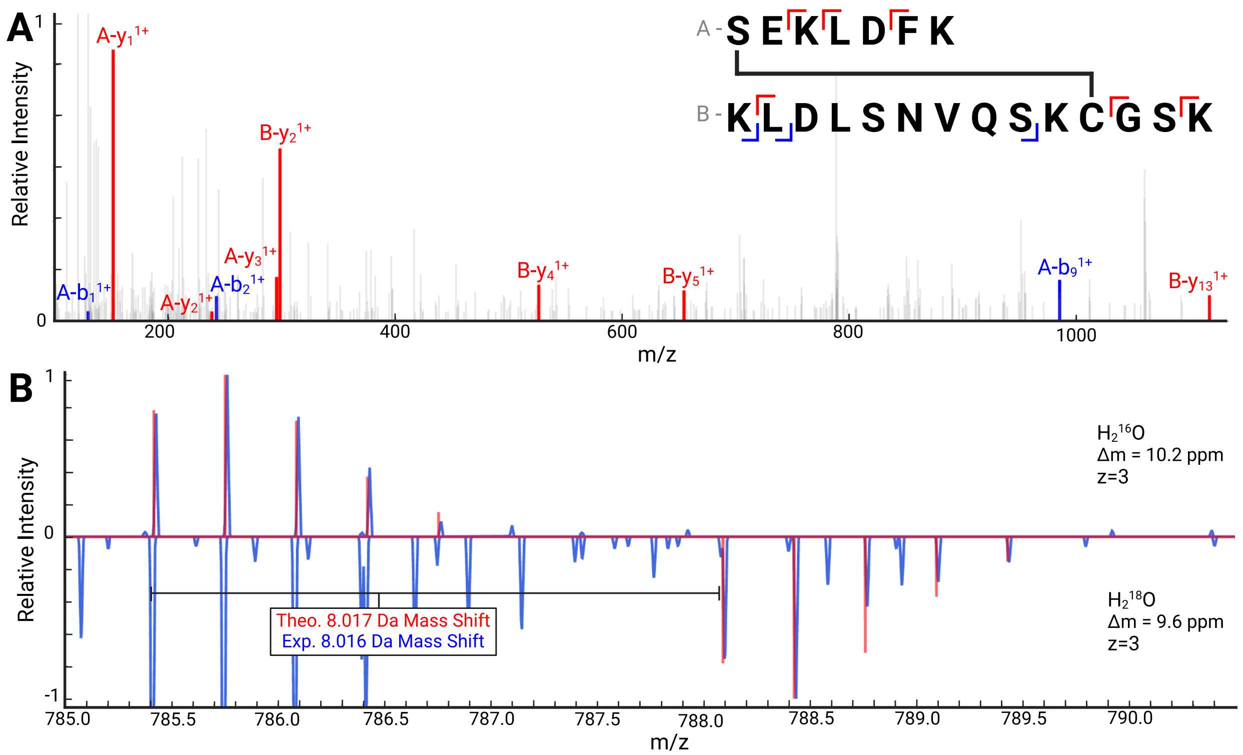


**Fig. S2: Evidence for a DHAA-mediated crosslink between residues S341-C291 in Tau**. **A**, The MS2 fragmentation spectrum for the precursor mass corresponding to the crosslinked species. It shows six y-ions and three b-ions, including one especially important fragment (B-$y_{13}$) that includes the crosslink. Fragmentation throughout the peptide is sufficient to accurately assign the crosslink to S341 but is ambiguous between K290 and C291. Other spectra localize the crosslink to C291 (data not shown). **B**, A mirror plot is shown with MS1 data derived from $H_2{}^{16}O$ samples on top (usual preparation) and $H_2{}^{18}O$ samples on the bottom (heavy isotope preparation). Raw data are shown in blue, and the theoretical isotope distributions are overlaid in red. The close match for each isotope peak compared to its theoretical value, for both mass and relative intensity, supports that the crosslink peptide has been identified correctly. Additional peaks in the $H_2{}^{18}O$ spectrum are from coeluting peptide species. $H_2{}^{18}O$ labeling validates the presence of a crosslink between two peptides containing residues S341 and C291 in Tau. Created with BioRender.com and Plotly.[34]

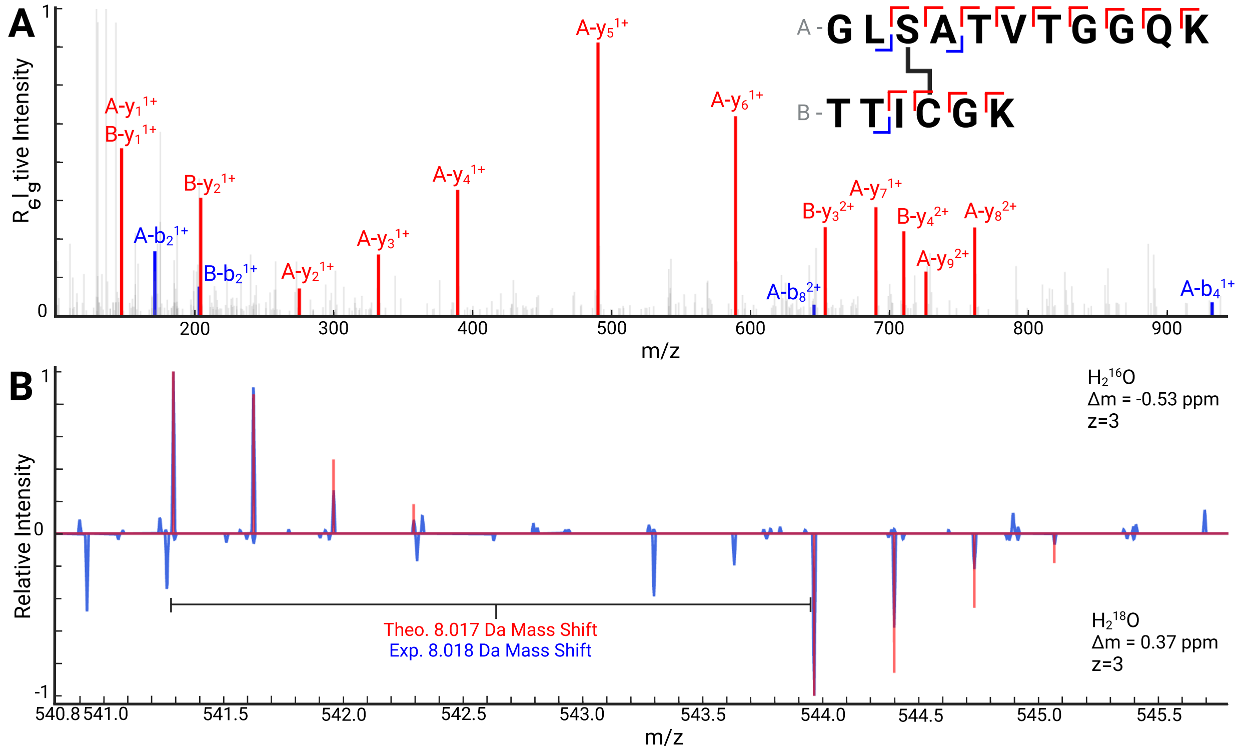


**Fig. S3: Evidence for a DHAA-mediated crosslink between residues S114-C109 in myelin proteolipid protein (PLP1)**. **A**, The MS2 fragmentation spectrum for the precursor mass corresponding to the crosslinked species. It shows thirteen y-ions and three b-ions, including four especially important fragments (A-$y_9$, A-$b_4$, B-$y_3$, and B-$y_4$) that include the crosslink. Fragmentation throughout the peptide is sufficient to accurately assign the crosslink to S114 and C109. **B**, A mirror plot is shown with MS1 data derived from $H_2{}^{16}O$ samples on top (usual preparation) and $H_2{}^{18}O$ samples on the bottom (heavy isotope preparation). Raw data are shown in blue, and the theoretical isotope distributions are overlaid in red. The match for each isotope peak compared to its theoretical value, for both mass and relative intensity, supports that the crosslink peptide has been identified correctly. Additional peaks in the $H_2{}^{18}O$ spectrum are from coeluting peptide species. $H_2{}^{18}O$ labeling validates the presence of a crosslink between two peptides containing residues S114 and C109 in PLP1. Created with BioRender.com and Plotly.[34]

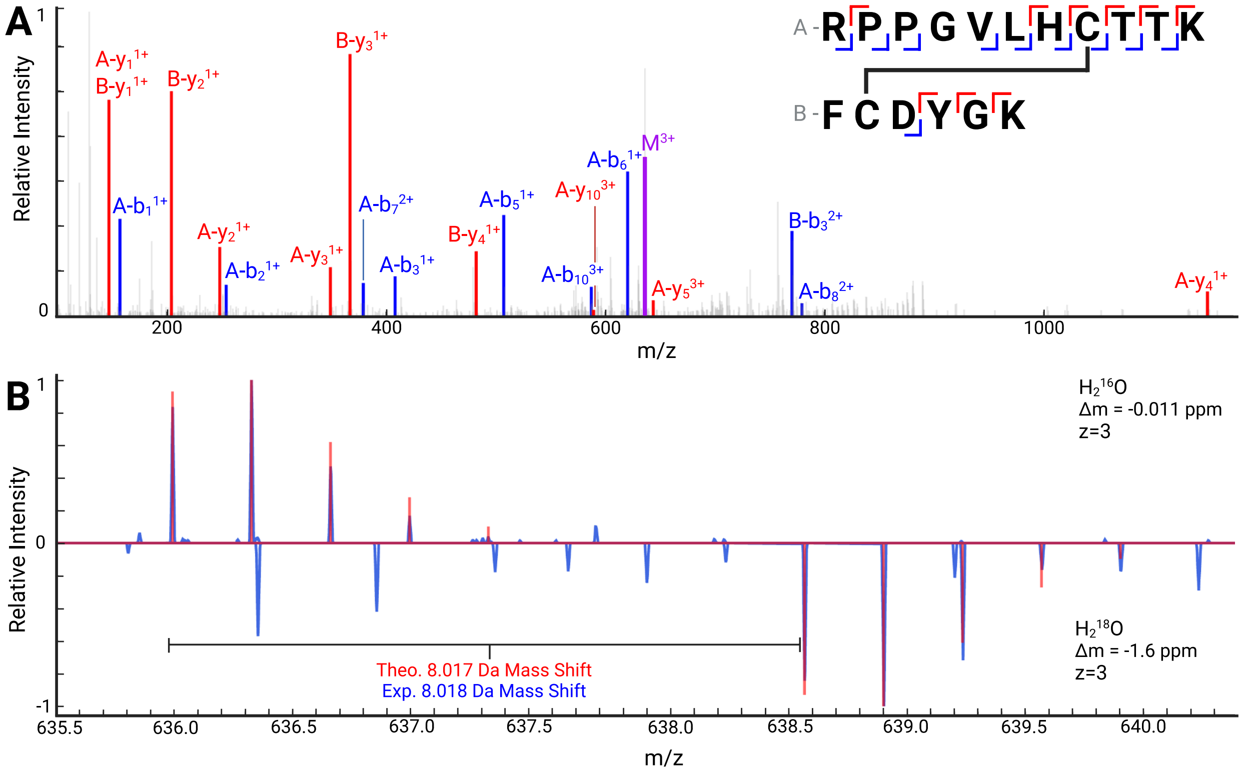


**Fig. S4: Evidence for a DHAA-mediated crosslink between residues C252-C257 in 2,′3′-cyclic nucleotide 3′-phosphodiesterase (CNP)**. **A**,The MS2 fragmentation spectrum for the precursor mass corresponding to the crosslinked species. It shows nine y-ions and ten b-ions, including seven especially important fragments (A-$y_4$, A-$y_5$, A-$y_{10}$, A-$b_8$, A-$b_9$, A-$b_{10}$, and B-$b_3$) that include the crosslink. A third type of ion ($M^{3+}$) is present, which is the unfragmented mass of the crosslinked peptides (shown in purple). Fragmentation throughout the peptide is sufficient to accurately assign the crosslink to C252 and C257. **B**, A mirror plot is shown with MS1 data derived from $H_2{}^{16}O$ samples on top (usual preparation) and $H_2{}^{18}O$ samples on the bottom (heavy isotope preparation). Raw data are shown in blue, and the theoretical isotope distributions are overlaid in red. The close match for each isotope peak compared to its theoretical value, for both mass and relative intensity, supports that the crosslink peptide has been identified correctly. Additional peaks in the $H_2{}^{18}O$ spectrum are from coeluting peptide species. $H_2{}^{18}O$ labeling validates the presence of a crosslink between two peptides containing residues C252 and C257 in CNP. Created with BioRender.com and Plotly.[34]

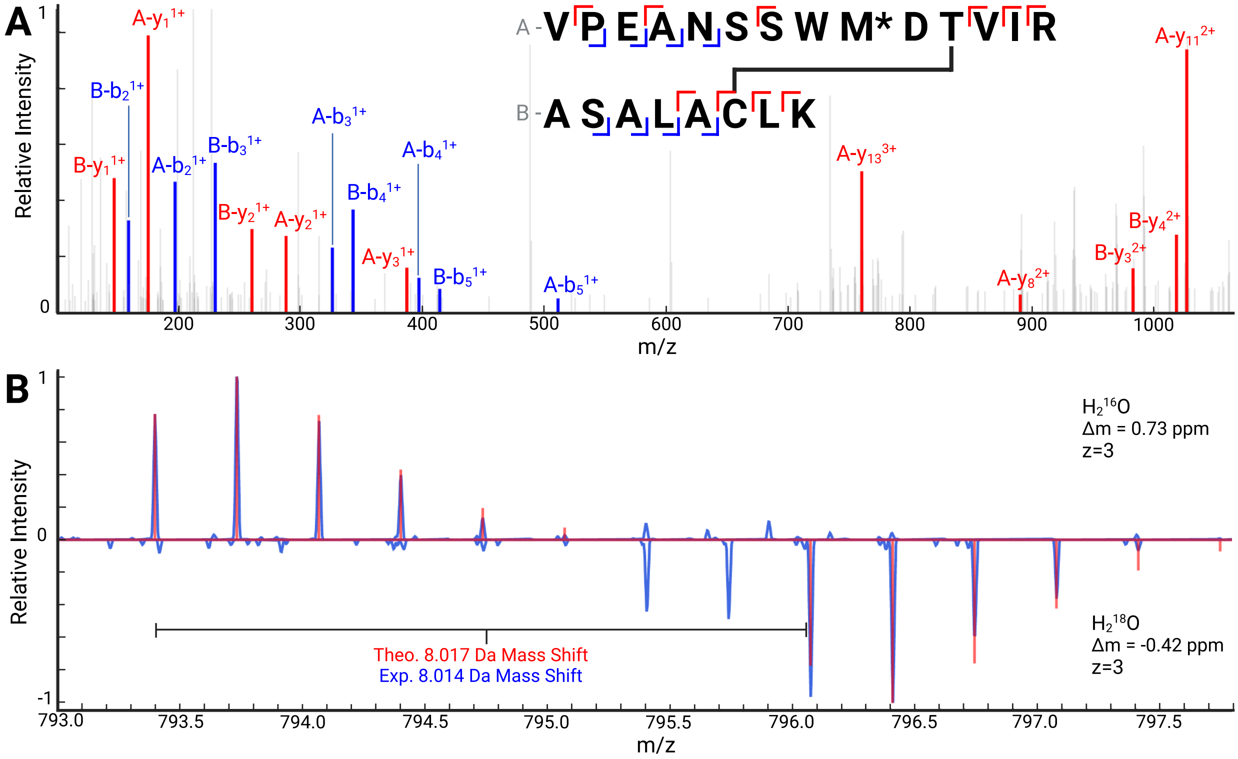


**Fig. S5: Evidence for a DHAA-mediated crosslink between residues S497 or T501 and C488 in dihydrolipoyl transacetylase (DLAT)**. **A**, The MS2 fragmentation spectrum for the precursor mass corresponding to the crosslinked species. "M*" represents that the methionine is oxidized. The MS2 spectrum shows ten y-ions and eight b-ions, including five especially important fragments (A-$y_8$, A-$y_{11}$, A-$y_{13}$, B-$y_3$, and B-$y_4$) that include the crosslink. Fragmentation throughout the peptide does not localize the crosslink to a particular DHAA residue, so it is ambiguous whether the crosslink is from S497 or T501 to C488. **B**, A mirror plot is shown with MS1 data derived from $H_2{}^{16}O$ samples on top (usual preparation) and $H_2{}^{18}O$ samples on the bottom (heavy isotope preparation). Raw data are shown in blue, and the theoretical isotope distributions are overlaid in red. The close match for each isotope peak compared to its theoretical value, for both mass and relative intensity, supports that the crosslink peptide has been identified correctly. Additional peaks in the $H_2{}^{18}O$ spectrum are from incomplete isotopic labeling of the C-terminal oxygens. $H_2^{18}O$ labeling validates the presence of a crosslink between two peptides containing residues S497/T501 and C488 in DLAT. Created with BioRender.com and Plotly.[34]

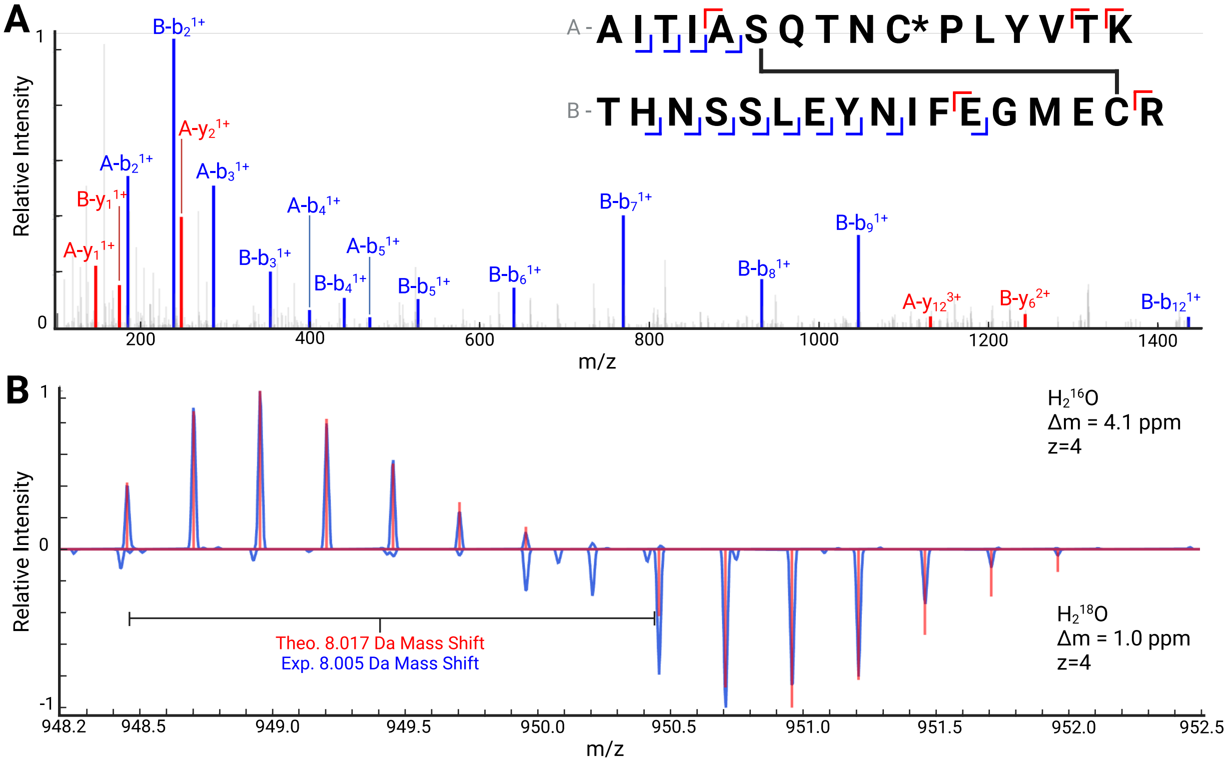


**Fig. S6: Evidence for a DHAA-mediated crosslink between residues S244 or T246 in collapsin response mediator protein 4 (CRMP4) and C439 in collapsin response mediator protein 2 (CRMP2)**. **A**, The MS2 fragmentation spectrum for the precursor mass corresponding to the crosslinked species. “C*” represents that the cysteine is carbamidomethylated. The MS2 spectrum shows five y-ions and thirteen b-ions, including two especially important fragments (A-$y_{12}$ and B-$y_6$) that include the crosslink. Fragmentation throughout the peptide does not localize the crosslink to a particular DHAA residue, so it is ambiguous whether the crosslink is from S244 or T246 to C439. **B**, A mirror plot is shown with MS1 data derived from $H_2{}^{16}O$ samples on top (usual preparation) and $H_2{}^{18}O$ samples on the bottom (heavy isotope preparation). Raw data are shown in blue, and the theoretical isotope distributions are overlaid in red. The close match for each isotope peak compared to its theoretical value, for both mass and relative intensity, supports that the crosslink peptide has been identified correctly. Additional peaks in the

$H_2{}^{18}O$ spectrum are from incomplete isotopic labeling of the C-terminal oxygens. $H_2{}^{18}O$ labeling validates the presence of a crosslink between two peptides containing residues S244/T246 in CRMP4 and C439 in CRMP2. Created with BioRender.com and Plotly.[34]

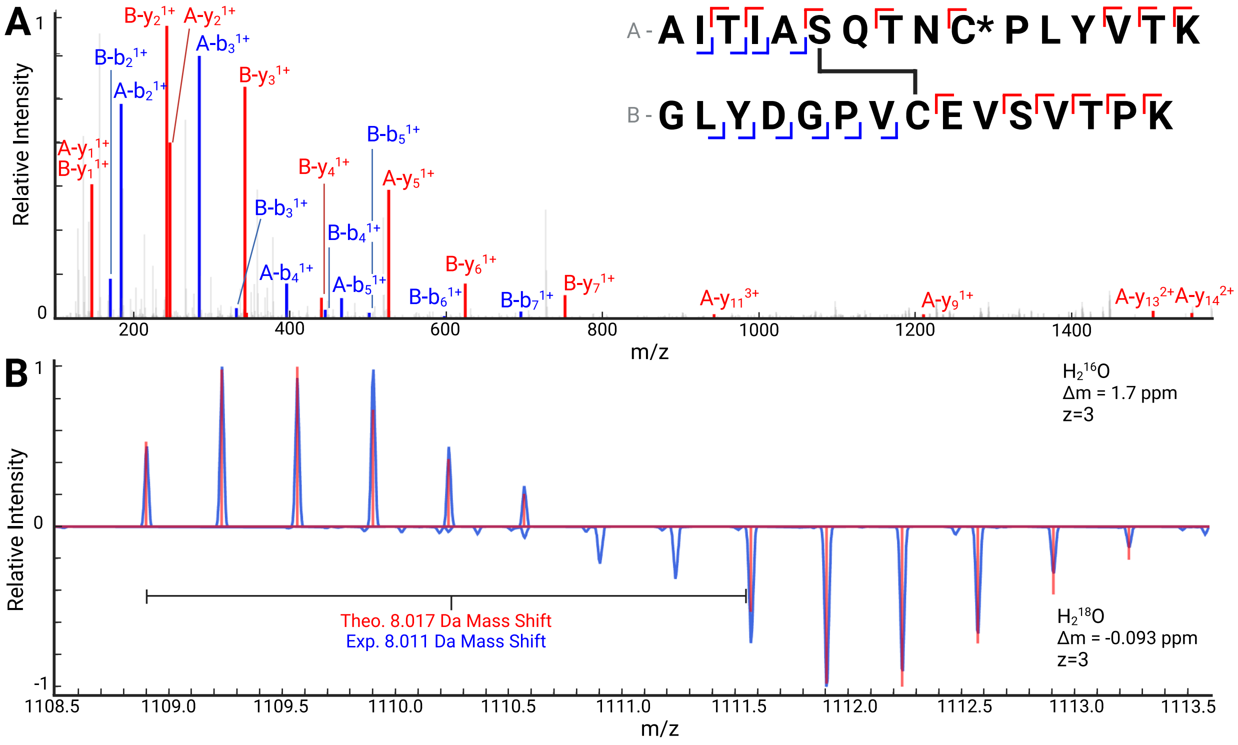


**Fig. S7: Evidence for a DHAA-mediated crosslink between residues S244 in collapsin response mediator protein 4 (CRMP4) and C504 in collapsin response mediator protein 2 (CRMP2)**. **A**, The MS2 fragmentation spectrum for the precursor mass corresponding to the crosslinked species. "C*" represents that the cysteine is carbamidomethylated. The MS2 spectrum shows fourteen y-ions and ten b-ions, including three especially important fragments (A-$y_{11}$, A-$y_{13}$, and A-$y_{14}$) that include the crosslink. Fragmentation throughout the peptide is sufficient to accurately assign the crosslink to S244 in CRMP4 and C504 in CRMP2. **B**, A mirror plot is shown with MS1 data derived from $H_2{}^{16}O$ samples on top (usual preparation) and $H_2{}^{18}O$ samples on the bottom (heavy isotope preparation). Raw data are shown in blue, and the theoretical isotope distributions are overlaid in red. The close match for each isotope peak compared to its theoretical value, for both mass and relative intensity, supports that the crosslink peptide has been identified correctly. Additional peaks in the $H_2{}^{18}O$ spectrum are from incomplete isotopic labeling of the C-terminal oxygens. $H_2{}^{18}O$ labeling validates the presence of a crosslink between two peptides containing residues S244 in CRMP4 and C504 in CRMP2. Created with BioRender.com and Plotly.[34]

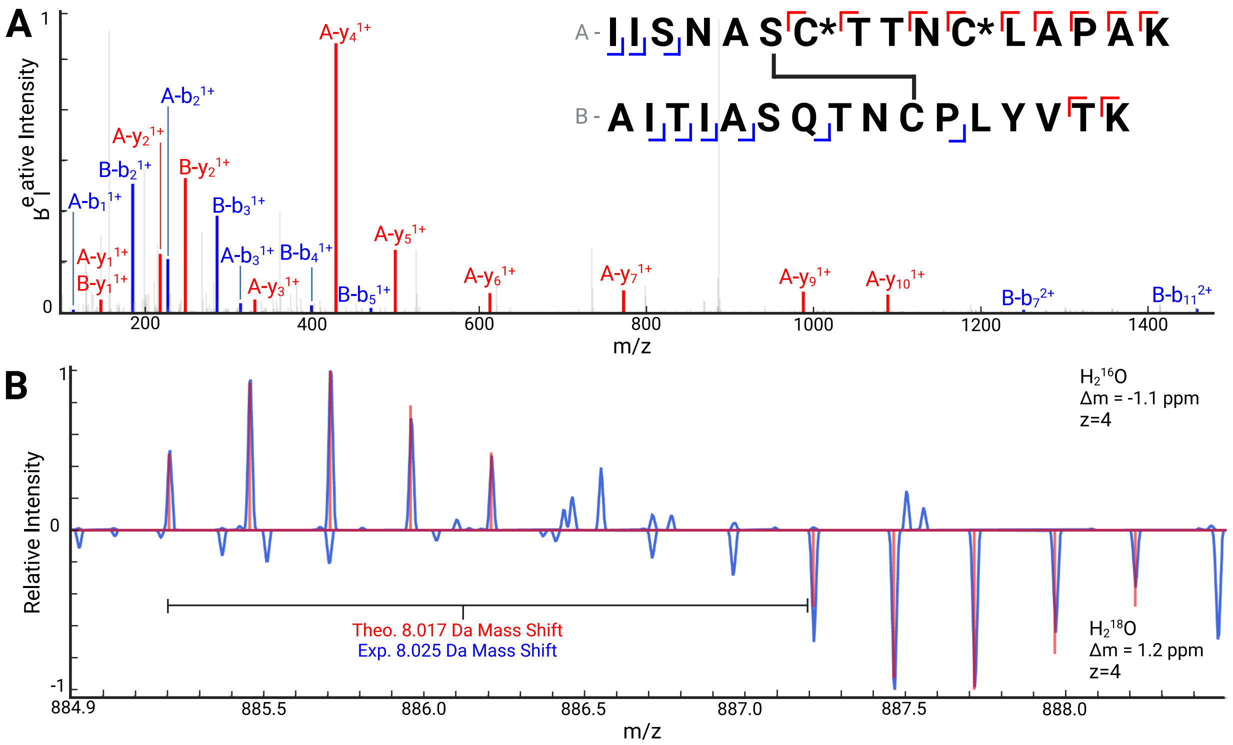


**Fig. S8: Evidence for a DHAA-mediated crosslink between residues S151 in glyceraldehyde-3-phosphate dehydrogenase (GAPDH) and C248 in collapsin response mediator protein 4 (CRMP4).** **A**, The MS2 fragmentation spectrum for the precursor mass corresponding to the crosslinked species. "C*" represents that the cysteine is carbamidomethylated. The MS2 shows eleven y-ions and nine b-ions, including one especially important fragment (B-$b_{11}$) that includes the crosslink. Fragmentation throughout the peptide is sufficient to accurately assign the crosslink to S151 in GAPDH and C248 in CRMP4. **B**, A mirror plot is shown with MS1 data derived from $H_2{}^{16}O$ samples on top (usual preparation) and $H_2{}^{18}O$ samples on the bottom (heavy isotope preparation). Raw data are shown in blue, and the theoretical isotope distributions are overlaid in red. The close match for each isotope peak compared to its theoretical value, for both mass and relative intensity, supports that the crosslink peptide has been identified correctly. Additional peaks in the $H_2{}^{18}O$ spectrum are from incomplete isotopic labeling of the C-terminal oxygens. $H_2{}^{18}O$ labeling validates the presence of a crosslink between two peptides containing residues S151 in GAPDH and C248 in CRMP4.

Created with BioRender.com and Plotly.[34]

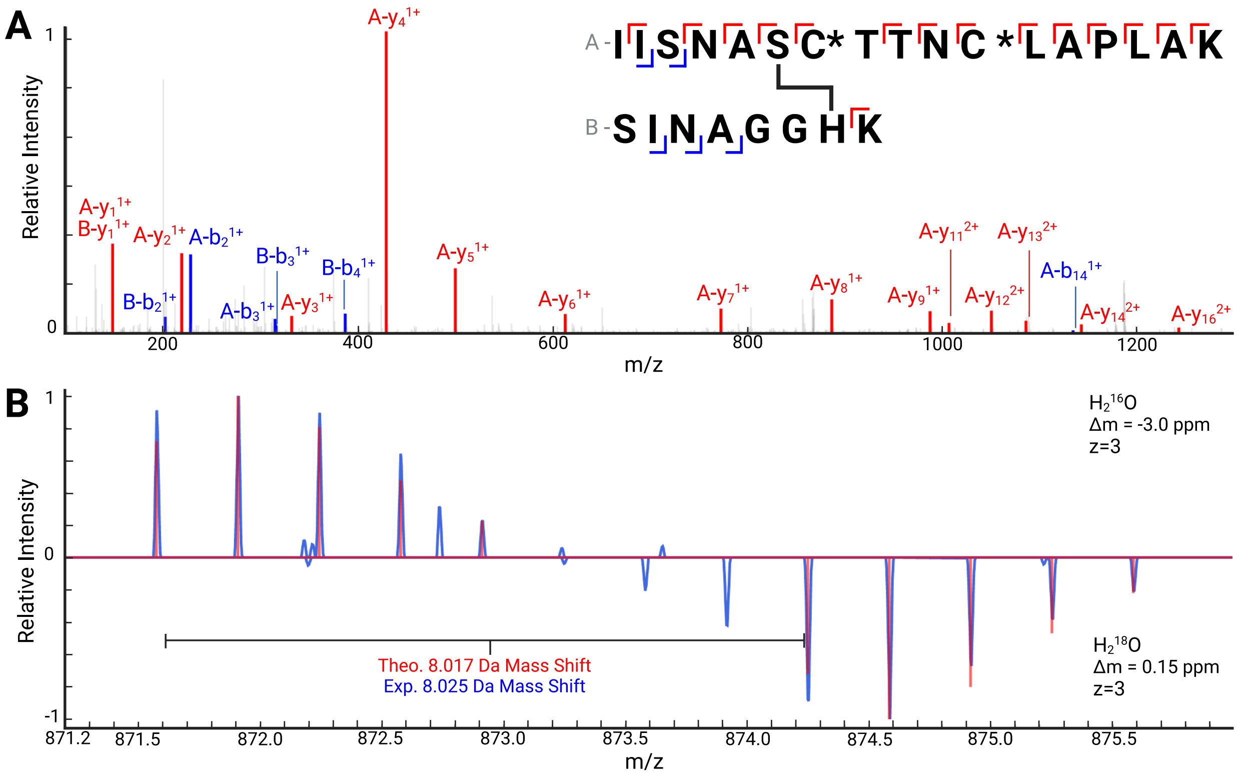


**Fig. S9: Evidence for a DHAA-mediated crosslink between residues S151 in glyceraldehyde-3-phosphate dehydrogenase (GAPDH) and H284 in voltage dependent anion channel 2 (VDAC2)**. **A**, The MS2 fragmentation spectrum for the precursor mass corresponding to the crosslinked species. "C*" represents that the cysteine is carbamidomethylated. The MS2 shows fifteen y-ions and five b-ions, including four especially important fragments (A-$y_{12}$, A-$y_{13}$, A-$y_{14}$, and A-$y_{16}$) that include the crosslink. Fragmentation throughout the peptide is sufficient to accurately assign the crosslink to S151 in GAPDH and H284 in VDAC2. **B**, A mirror plot is shown with MS1 data derived from $H_2{}^{16}O$ samples on top (usual preparation) and $H_2{}^{18}O$ samples on the bottom (heavy isotope preparation). Raw data are shown in blue, and the theoretical isotope distributions are overlaid in red. The close match for each isotope peak compared to its theoretical value, for both mass and relative intensity, supports that the crosslink peptide has been identified correctly. Additional peaks in the $H_2{}^{18}O$ spectrum are from incomplete isotopic labeling of the C-terminal oxygens. $H_2{}^{18}O$ labeling validates the presence of a crosslink between two peptides containing residues S151 in GAPDH and H284 in VDAC2. Created with BioRender.com and Plotly.[34]

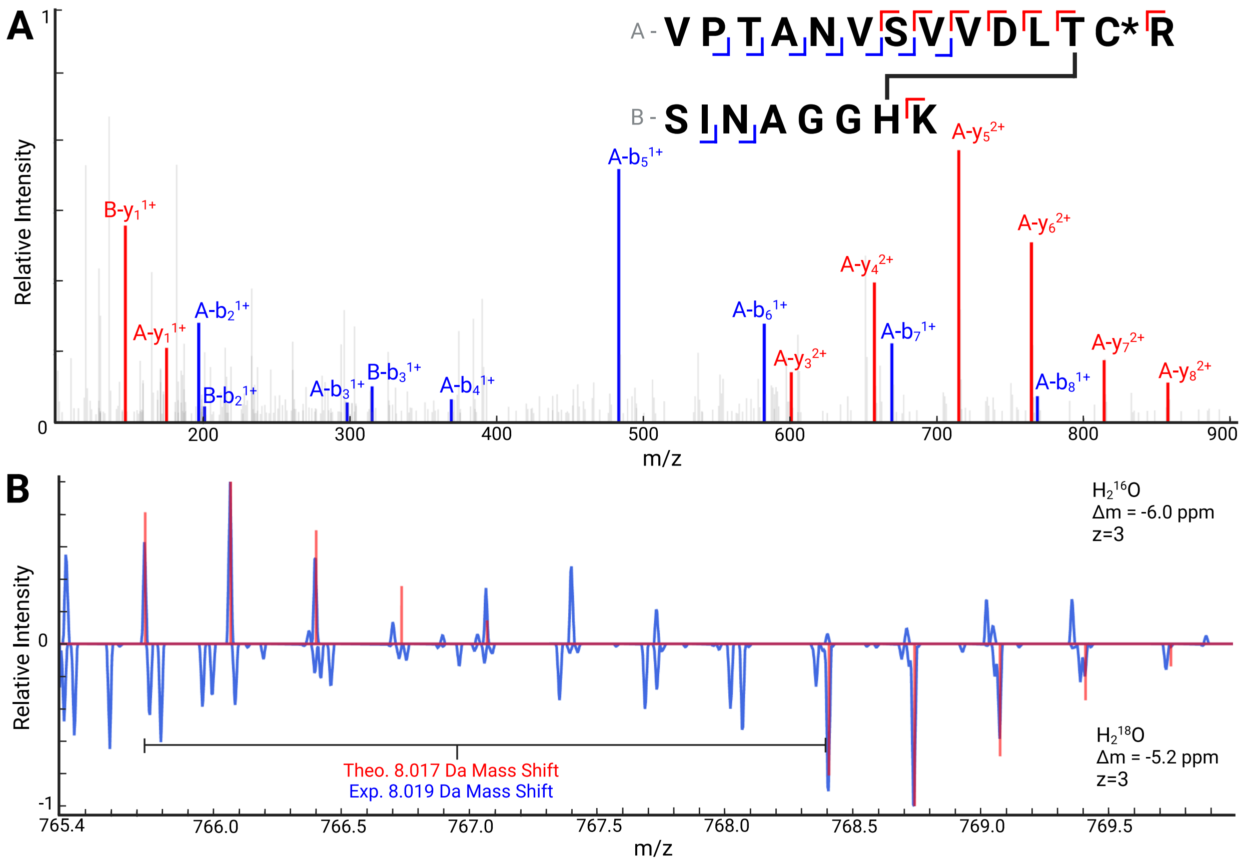


**Fig. S10: Evidence for a DHAA-mediated crosslink between residues T246 in glyceraldehyde-3-phosphate dehydrogenase (GAPDH) and H284 in voltage dependent anion channel 2 (VDAC2)**. **A**, The MS2 fragmentation spectrum for the precursor mass corresponding to the crosslinked species. "C*" represents that the cysteine is carbamidomethylated. The MS2 shows eight y-ions and nine b-ions, including six especially important fragments (A-$y_3$, A-$y_4$, A-$y_5$, A-$y_6$, A-$y_7$, and A-$y_8$) that include the crosslink. Fragmentation throughout the peptide is sufficient to accurately assign the crosslink to T247 in GAPDH and H284 in VDAC2. **B**, A mirror plot is shown with MS1 data derived from $H_2{}^{16}O$ samples on top (usual preparation) and $H_2{}^{18}O$ samples on the bottom (heavy isotope preparation). Raw data are shown in blue, and the theoretical isotope distributions are overlaid in red. The match for each isotope peak compared to its theoretical value, for both mass and relative intensity, supports that the crosslink peptide has been identified correctly. Additional peaks in the $H_2{}^{18}O$ spectrum are from incomplete isotopic labeling of the C-terminal oxygens and coeluting peptide species. $H_2{}^{18}O$ labeling validates the presence of a crosslink between two peptides containing residues T247 in GAPDH and H284 in VDAC2. Created with BioRender.com and Plotly.[34]

**Fig. S11: Relative occupancy of modifications at each protein residue that has been identified as an eliminylation site.** The legend defining colors for the various modifications is shown next to the first plot and pertains to all plots. Occupancy was estimated for each eliminylation site that was (i) identified in the Steen sarkosyl-insoluble datasets; (ii) identified by a DHAA conjugate as a site of eliminylation; and (iii) found in at least 50% of biological replicates.

All data presented here are occupancy estimates based on FlashLFQ normalized intensities. These data are extrapolated from mass spectral intensities and are complex measurements that are ultimately determined by several factors, such as peptide abundance, coeluting species abundance, and ionization efficiencies of each species. The raw data used to assemble these graphs is available in Table S4.

Each eliminylation site residue is listed along the x-axis. Relative occupancy fraction (y-axis) is displayed as stacked bars for the various modifications. Data is derived from the sarkosyl-insoluble samples from the Steen dataset for AD specimens (left bar) and age-matched controls (right bar) for each residue. Error bars represent the standard error of the mean. Figures created with Plotly.[34]

**Tau Occupancy** **Legend**

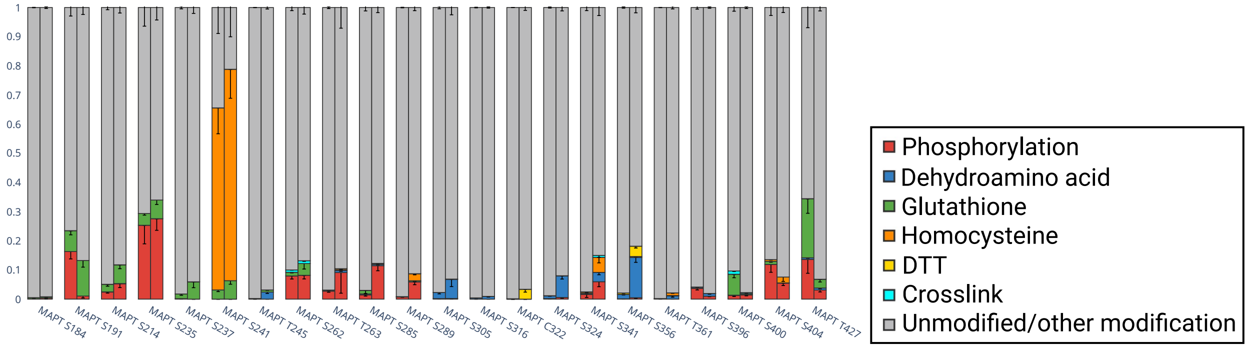

#### CRMP2 Occupancy

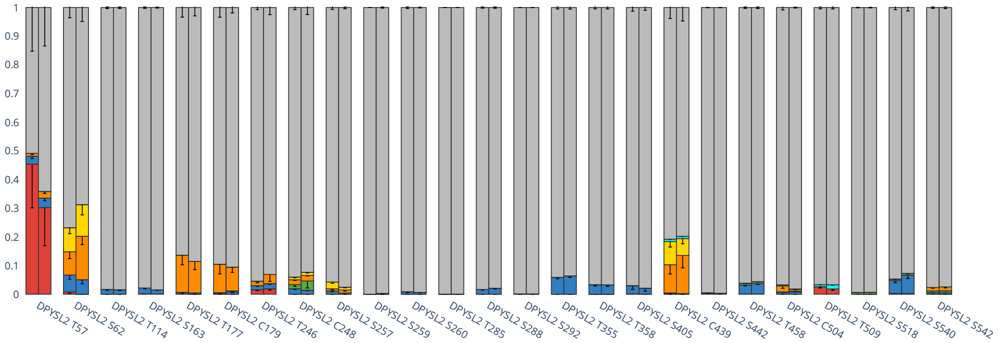


#### GAPDH Occupancy

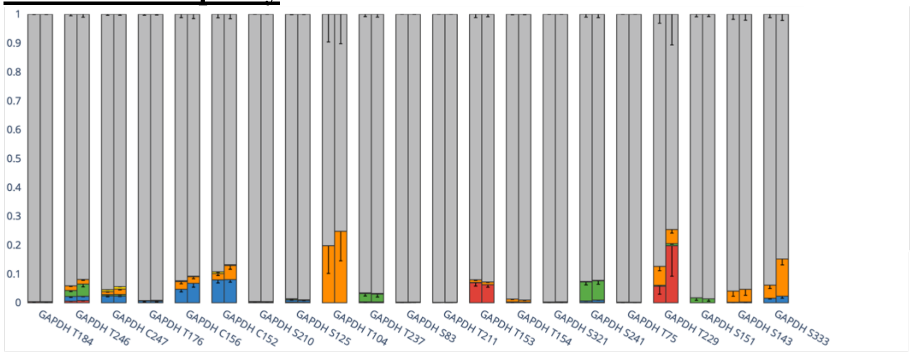


#### PKM Occupancy

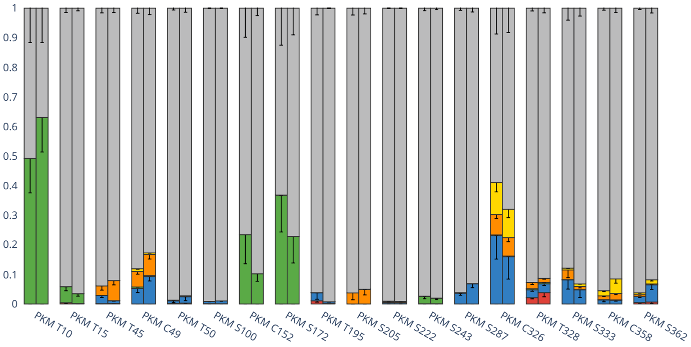


#### VCAN Occupancy

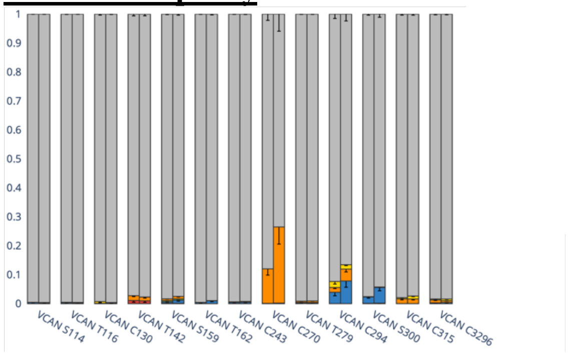

### All Other Proteins

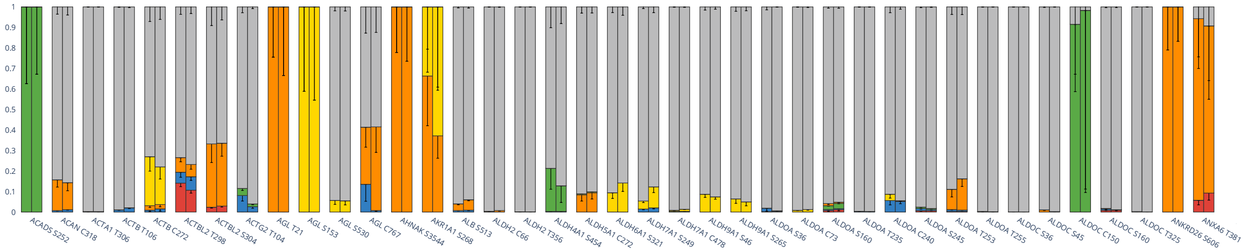


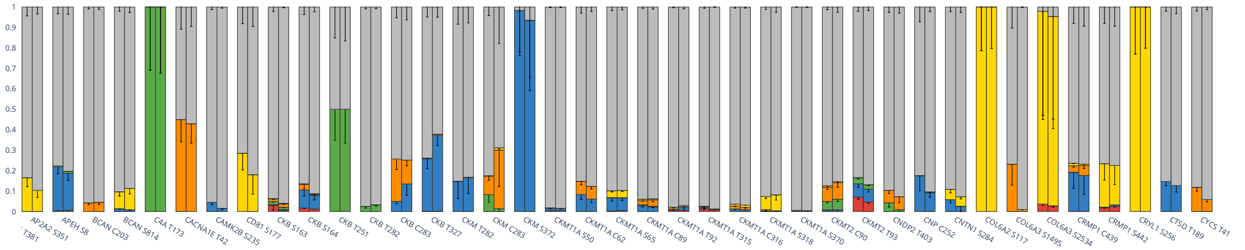


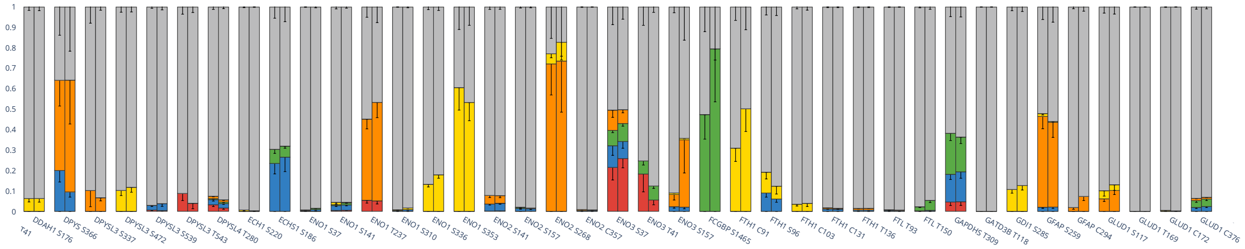


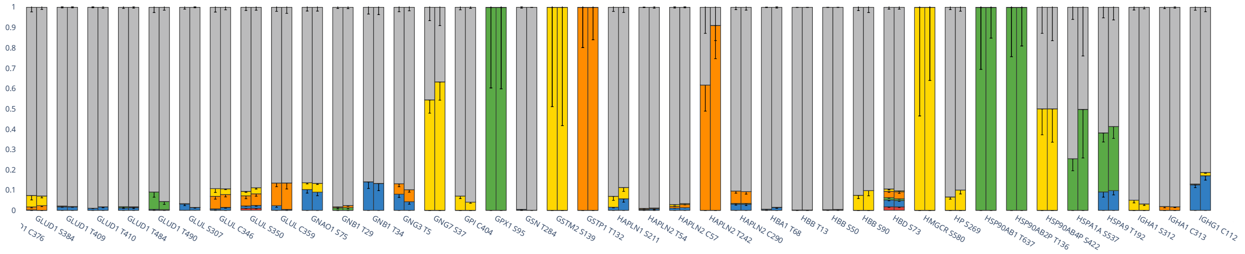


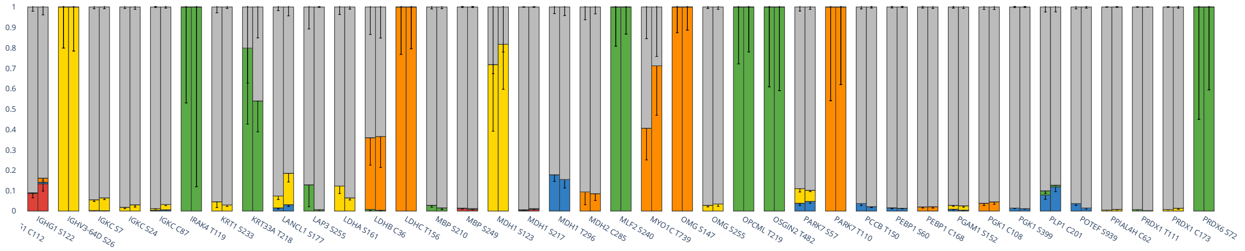

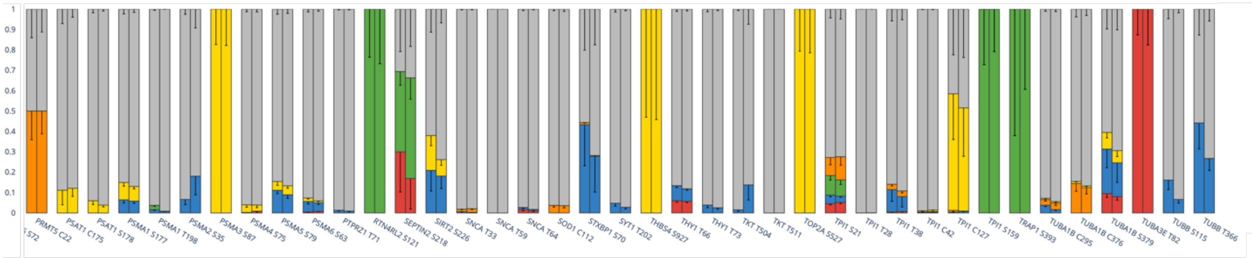

1
0.9
0.8
0.7
0.6
0.5
0.4
0.3
0.2
0.1
0
PRMT5 C22
PSAT1 C175
PSAT1 S178
PSMA1 S177
PSMA1 T198
PSMA2 S35
PSMA3 S87
PSMA4 S75
PSMA5 S79
PSMA6 S63
PTPRZ1 T71
RTN4RL2 S121
SEPTIN2 S218
SIRT2 S226
SNCA T33
SNCA T59
SNCA T64
SOD1 C112
STXBP1 S70
SYT1 T202
THBS4 S927
THY1 T66
THY1 T73
TKT T504
TKT T511
TOP2A S527
TPI1 S21
TPI1 T28
TPI1 T38
TPI1 C42
TPI1 C127
TPI1 S159
TRAP1 S393
TUBA1B C295
TUBA1B C376
TUBA1B S379
TUBA3E T82
TUBB S115
TUBB T366


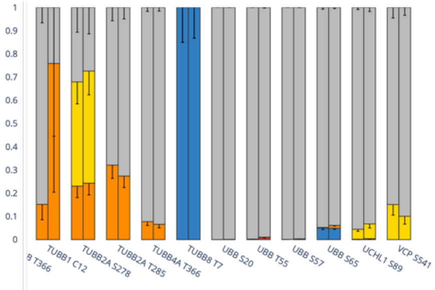

1
0.9
0.8
0.7
0.6
0.5
0.4
0.3
0.2
0.1
0
TUBB1 C12
TUBB2A S278
TUBB2A T285
TUBB4A T366
TUBB8 T7
UBB S20
UBB T55
UBB S57
UBB S65
UCHL1 S89
VCP S541

**Table S1:** Brain specimen details.

| Sample | Condition | Age | Sex | Post-mortem interval (hh:mm) |
|---|---|---|---|---|
| BA339 | AD | 87 | F | 3:56 |
| BA359 | AD | >90 | M | 4:40 |
| BA389 | AD | 76 | M | 18:03 |
| BA406 | AD | 85 | M | 14:16 |
| BA408 | AD | 80 | Unknown | 14:05 |
| BA420 | AD | 89 | Unknown | 4:15 |
| BA432 | AD | 87 | Unknown | 11:45 |
| BA483 | AD | >90 | Unknown | 24:35 |
| BA500 | AD | 80 | Unknown | 6:00 |
| BA533 | AD | >90 | M | 23:39 |
| BA130 | Control | 81 | F | 19:45 |
| BA186 | Control | 55 | F | 4:27 |
| BA452 | Control | 70 | F | 5:05 |

**Table S2:** Collection times (min.) for the eight fractions obtained from offline fractionation of tryptic peptide samples.

| | Pool 1 | Pool 2 |
|---|---|---|
| Fraction 1 | 0-4 | 19-20.5 |
| Fraction 2 | 4-7 | 20.5-22 |
| Fraction 3 | 7-10 | 22-23.5 |
| Fraction 4 | 10-13 | 23.5-25 |
| Fraction 5 | 13-14 | 25-28 |
| Fraction 6 | 14-15 | 28-31 |
| Fraction 7 | 15-17 | 31-35 |
| Fraction 8 | 17-19 | 35-40 |

**Tables S3 and S4 can be found in Supplementary Materials 4 and 5 respectively.**

**Table S5: Numbers of eliminylation sites and proteins containing DHAAs and DHAA conjugates.** This table was generated with data exclusively from Wesseling *et al.* (Steen data) to support that DHAAs are an order of magnitude more abundant in sarkosyl-insoluble samples. The comparison in the main text combined identifications made from both Steen and Smith data. This table shows results of a similar comparison of unfractionated samples for both sarkosyl-insoluble and sarkosyl-soluble sample types, yielding a similar order-of-magnitude difference in eliminylation sites.

| | **Sarkosyl-Insoluble** | | **Sarkosyl-Soluble** | |
|---|---|---|---|---|
| | Sites | Proteins | Sites | Proteins |
| Any Identification | 489 | 195 | 22 | 6 |
| >50% of Specimens | 93 | 269 | 11 | 18 |
| >80% of Specimens | 44 | 100 | 8 | 8 |
| 100% of Specimens | 7 | 8 | 0 | 0 |

**Table S6:** Crosslinks potentially identified, but not included in the main list of 11 crosslinks (Table 3) due to missing some aspect(s) of strong evidence. "Weak MS2" refers to the MS2 spectra for these species not being strongly supportive (contained insufficient fragments and did not show evidence of fragments with the crosslink). "1% FDR" means the species was not identified in a search against the whole proteome as a modification at the site of interest, as described in "*Final bottom-up proteomics search with crosslink-appended, GPTMD-modified Swiss-Prot database*." in Methods.

| Protein | DHA Residue | Protein | Nucleophile | Reason for Rejection |
|---|---|---|---|---|
| Amyloid-β | S26 | Amyloid-β | H6 | $H_2{}^{18}O$ labeling did not validate |
| NEFM | T264 | FGG | H426 | Weak MS2, 1% FDR |
| NEFM | S592 | CRMP2 | K254 | Weak MS2, 1% FDR |
| CRMP4 | T246 | PLP | C228 | Missed Monoisotopic |
| CRMP4 | T246 | GAPDH | C152 | Weak MS2, 1% FDR |
| CRMP4 | T246 | GAPDH | C247 | Weak MS2, 1% FDR |
| ENO3 | C399 | ENO3 | H190\|H191 | Weak MS2, 1% FDR |
| ENO3 | C357 | ENO3 | H371 | Weak MS2, 1% FDR |
| ENO2 | S370 | ENO2 | C357 | Weak MS2, 1% FDR |
| Midkine | C94 | Midkine | C126 | 1% FDR search |

**Tables S7 and S8 can be found in Supplementary Materials 6 and 7 respectively.**

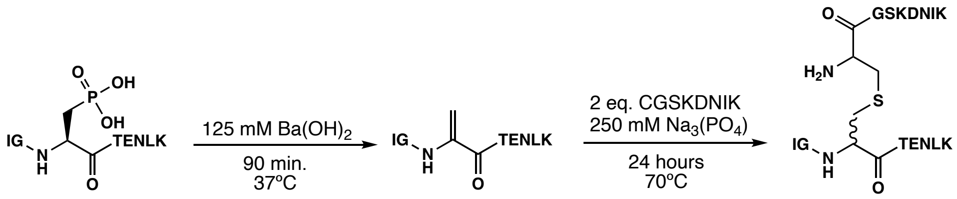


**Scheme S1:** Eliminylation and subsequent crosslinking reaction to generate the S262-C291 synthetic standard from model peptides.

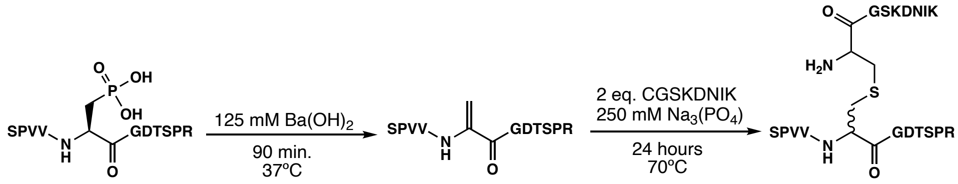


**Scheme S2:** Eliminylation and subsequent crosslinking reaction to generate the S400-C291 synthetic standard from model peptides.
